# Programmable Mixed-Signal Biocomputers in Mammalian Cells

**DOI:** 10.1101/2022.06.07.495130

**Authors:** Justin H. Letendre, Benjamin H. Weinberg, Marisa Mendes, Jeffery M. Marano, K. J. William Benman, Rachel Petherbridge, Kamila Drezek, Samantha E. Koplik, Alexandra Piñeiro, Wilson W. Wong

**Author notes:** Correspondence and requests for materials should be sent to this author.

## Abstract

Living cells perform sophisticated computations that guide them toward discrete states. Synthetic genetic circuits are powerful tools for programing these computations, where transcription-regulatory networks and DNA recombination are the two dominant paradigms for implementing these systems. While each strategy exhibits unique strengths and weaknesses, integrating both into one seamless design framework would enable advanced gene circuit designs intractable with either approach alone. Here, we present Computation via Recombinase Assisted Transcriptional Effectors (CREATE), which leverages site-specific recombination to perform robust logic on discreet computational layers and programmable transcription factors that connect these layers, allowing individual calculations to contribute toward larger operations. We demonstrate the functionality of CREATE by producing sophisticated circuits using a simple plug-and-play framework, including 189 2-input-3-output circuits, modular digital-to-analog signal converters, a 2-bit multiplier circuit, and a digital and analog mixed-signal generator. This work establishes CREATE as a versatile platform for programming complex signal processing systems capable of high-fidelity logic computation and tunable control over circuit output levels.

**One-Sentence Summary:** We present a minimal and robust genetic circuit platform for programming cells with sophisticated signal processing capabilities.

## Main Text

Endogenous gene circuits are adept at processing combinations of input signals to produce different types and levels of gene expression outputs. This signal processing is vital for cells to interact with their environment and coordinate responses [1]. Synthetic versions of these gene circuits allow for precise control over living systems by programming cells with custom signal processing and computational abilities.

Since the introduction of synthetic genetic circuits in the early 2000’s [2–9] [10, 11] [12–17], two prominent design paradigms have emerged: transcription/post-transcription networks that mimic natural systems and recombinase/nuclease-based systems that rearrange DNA to perform computations. Transcription factors (TFs) and regulatory RNAs are adept at programming analog circuits and can be used to tailor the transcriptional output from a target promoter (**Figure 1A, left**). While progress has been made in prokaryotic systems [7] [18], the ability to modulate gene expression levels in response to large sets of user- or environmentally-defined cues in mammalian cells remains lacking. Those that have been generated demonstrate high functionality rely on ad hoc designs that are not easily reconfigurable and would be difficult to scale up [9, 11] [17].

**Figure 1:**
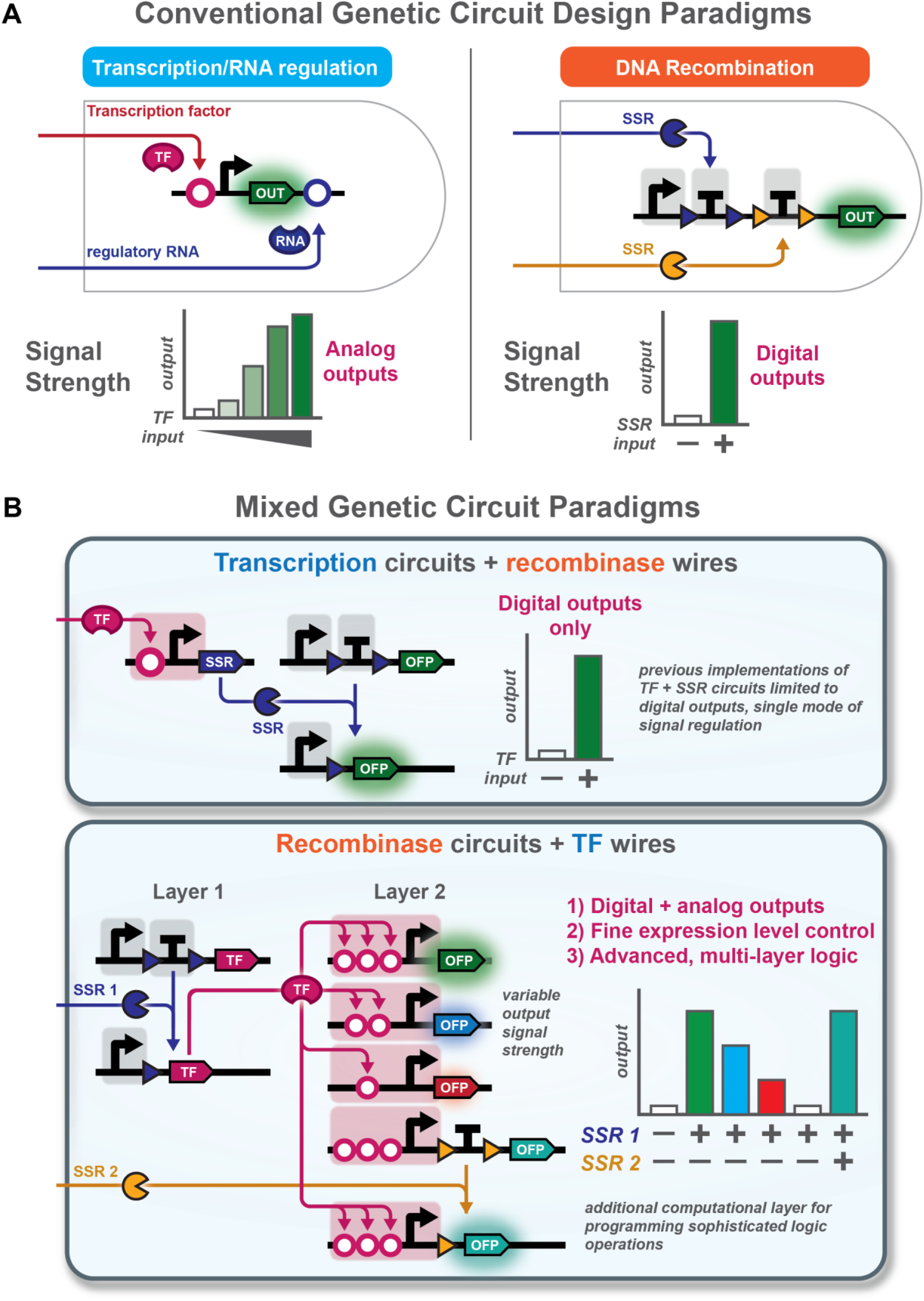
SSRs driving TF production expands computational power and allows both analog and digital computation. **A**) Two of the most prevalent paradigms in genetic circuit designs are either transcription factor (TF) and regulatory RNA-based networks (left) or DNA recombination-based using site-specific recombinases (SSRs) and nucleases (right). TF networks are adept at programming analog signaling responses where TFs target engineered promoters to produce a specific level of output signal but have been traditionally difficult to use for programming complex logic computation. Conversely, SSRs are adept at performing logic operations through DNA excision reactions such as removing a terminator sequence to initiate transcription of an output, as shown here, but typically yield digital outputs with no capacity for analog output signals **B**) Circuits employing both TFs and SSRs have previously used TFs to drive SSR expression (top panel), useful for signal amplification but limiting in the regulation of multiple output genes and the strength of the output signals. By placing TFs in a circuit under the control of SSR inputs (bottom panel), different TFs can be selected to target (1) multiple output signals, (2) promoters of varying transcriptional strength for differential signal intensity, and (3) other circuits receiving further SSR input signals for complex signal processing and logic computation that is relatively easy to program.

Unlike TF/RNA networks, site-specific recombinases (SSRs) are uniquely capable for performing digital logic operations in both prokaryotic and eukaryotic systems, including mammalian cells [4, 5] [8, 18] (**Figure 1A, right**). Recently, the BLADE circuit design has demonstrated a flexible approach for creating complex yet robust logic circuits capable of integrating up to 6 unique input signals, including a field-programmable logic lookup table and multiple arithmetic tools [5]. While SSR-based systems are powerful for developing genetic logic circuits, they lack a fundamental capability of signal processing systems: they cannot regulate their level of output response, only whether it is present or absent. This precludes their use in settings that require precise engineering of output signals, such as integration of multiple external stimuli to generate differential response levels [19].

Previous attempts to combine the analog programmability of TFs and regulatory RNAs with SSR-mediated digital logic and memory have focused on placing SSRs under the control of a TF [18, 20] [21–24] (**Figure 1B, top**). While this design allows for potent signal amplification and information transfer with a high signal-to-noise ratio, it is fundamentally limiting in scope for the types of computation that may be performed. In contrast, if the roles were reversed and SSR-based circuits were used to control TF expression, multiple computational layers could be wired together similar to the function of a computer chip comprised of individual circuits (**Figure 1B, bottom**). This alternative design paradigm would provide several advantages over more classic circuit design approaches, such as the ability to reuse computational parts in multiple circuit layers and allow for complex signal processing operations to be performed on either a limited or broad range of inputs.

To achieve this, we present an integrated genetic circuit design platform consists of recombinase circuits and programmable transcription factors for 1) performing combinatorial input signal sensing to specify output gene expression levels and 2) assembling multiple lower-order gene circuits to perform computations previously intractable for biological implementation. This multi-layered circuit design, <u>C</u>omputation via <u>R</u>ecombinase <u>A</u>ssisted <u>T</u>ranscriptional <u>E</u>ffectors (CREATE), operates as a two-layer cascade that connects multiple levels of BLADE circuits using programmable TFs (**Figure 2A**). Given a unique set of SSR inputs, desired TFs are selected from Layer 1 and target engineered promoters responsive to each TF in Layer 2. To maximize its functionality, we have designed the CREATE platform to interface with both CRISPRa TFs and synthetic zinc finger transcriptional regulators (synZiFTRs). We demonstrate the capabilities of this platform by rapidly prototyping 189 unique 2-input-3-output circuits using modular components, of which 90% compute their intended truth tables without the need for optimization. We further leverage CREATE to assemble multiple digital-to-analog circuits (DACs) using a library of custom promoters, and demonstrate differential regulation of multiple protein expression levels in the same cell. Additionally, we present a genetic 2-bit multiplier circuit, which computes the product of up to four SSR inputs in mammalian cells, and the first biological device capable of dual analog and digital signal generation. This work presents a highly reconfigurable platform for assembling some of the most advanced biological signal processing systems to date. CREATE provides an adaptable framework for coupling digital logic processing and memory with analog output signals, similar to the function of native gene regulatory and cell-cell communication networks.

**Figure 2:**
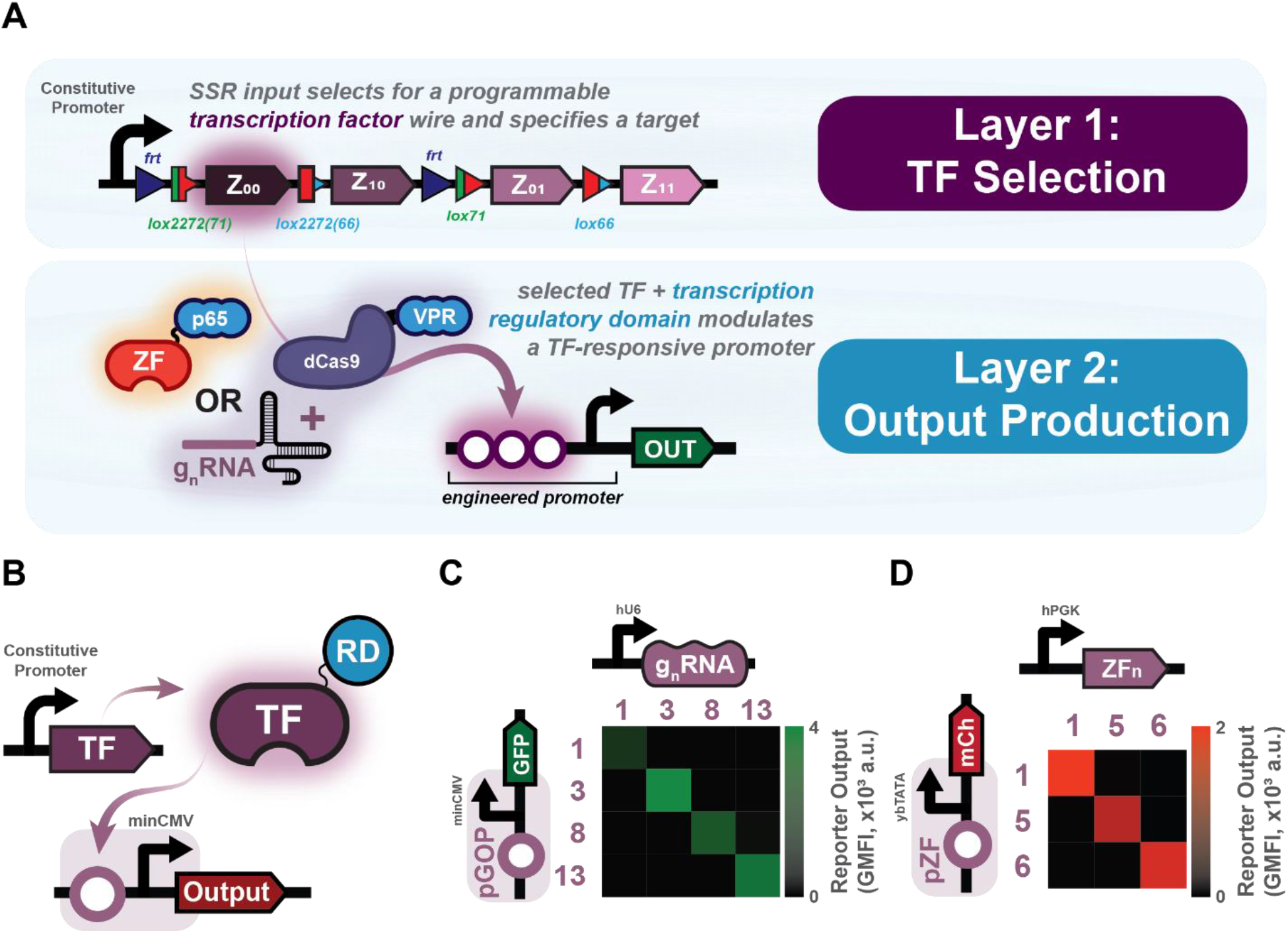
The CREATE platform combines recombinase logic circuits with programmable transcription factor ‘wires’. **A**) The multi-layer circuit design is connected by programmable transcription factors (TFs). TFs are chosen by recombinase inputs to a Layer 1 TF Decoder circuit, and target cognate promoters in Layer 2 to produce a desired level of output. **B**) Several orthogonal gRNAs and synZiFTRs have been validated for use with the Layer 1 circuit. Each TF was transfected with either a cognate or off-target promoter and the mean fluorescence output signal is plotted for each interaction. Heat maps for gRNA (**C**) and synZiFTR (**D**) candidates represent the geometric mean of flow cytometry data from three technical replicates (*n = 3*) of transiently transfected HEK293FT cells collected 48 hours post transfection.

## Results

### Validating Orthogonal transcriptional activators

The CREATE platform is composed of two circuit layers, each based on the BLADE single-layer circuit template [5]. A transcription factor (TF) is selected for using up to two site-specific recombinase (SSR) inputs (Cre and Flp) in Layer 1 and connects to a cognate promoter in Layer 2 to produce a unique transcriptional output (**Figure 2A**). Each individual layer performs its own computation, similar to integrated circuit chips in an electronic circuit. Each ‘chip’ is connected by TF ‘wires’ to produce a user-defined function.

Designing an effective layered circuit requires a library of orthogonal TFs to use as wiring. We built an *in silico* library of >1000 randomly generated guide-RNA (gRNA) spacer sequences and processed them using the CRISPR Optimized Design tool [25] to ensure the orthogonality of each candidate against potential target sites in the human genome. 18 of the gRNA spacer sequences had an orthogonality score ≥98% and were cloned into a gRNA scaffold template to test (**Table S1**). Cognate gRNA-responsive promoters were generated by placing a single operator site complementary to the gRNA immediately upstream of a minimal CMV promoter (**Table S1**). When paired with the CRISPRa dCas9-VPR activator in human embryonic kidney cells (HEK293FT), each gRNA candidate yields reporter output from its target promoter with a high dynamic range up to 37x (**Figure S1**).

We validated the orthogonality of four gRNA candidates displaying the highest dynamic ranges by transfecting them with each gRNA-responsive promoter, observing minimal off-target effects of any gRNA acting on non-cognate operator sites (**Figure 2B, C**). To expand the programmability of our platform and demonstrate its utility using human-derived TFs, we also validated the orthogonality of three synZiFTRs that have previously demonstrated high performance in mammalian cells *in vitro* and *in vivo* [26]. Each synZiFTR is composed of a zinc finger binding domain conjugated to a p65 activation domain and displays both high dynamic range and orthogonality to one another in HEK293FT cells (**Figure 2B,D**).

### Optimizing SSR target sites and off-target expression to promote efficient multi-layer communication

When placing the gRNAs into an un-optimized BLADE circuit design we find that the output from gRNA reporters was relatively weak compared with constitutively expressing gRNAs (data not shown). We hypothesize that SSRs binding to their target sites after circuit cleavage may sterically hinder pol III binding to the hU6 promoter, thus decreasing the transcription rate of gRNAs (**Figure 3A**). To alleviate this constraint, we systematically varied the spacing between the hU6 transcription start site (TSS) driving gRNA production and a Cre recombinase target site (loxP) upstream of a gRNA sequence to determine the optimal spacing requirements to promote more efficient transcription. When the loxP-gRNA constructs are transfected in HEK293FT cells without Cre recombinase, reporter output decreases monotonically with increasing space between the TSS and loxP-gRNA sequence (**Figure 3B**). However, when Cre is transfected with the gRNA and reporter plasmids, we note increasing gRNA output up to 100 bp spacing between the TSS and loxP target, followed by decreasing reporter expression at higher 5’ spacer lengths (**Figure 3B**). However, gRNA reporter expression at 100 bp spacing between the TSS and gRNA sequence is only ∼75% of the output from samples without SSR addition. Two possibilities could account for this: 1) that SSR binding to its target site after cleavage interrupts transcription by blocking the polymerase from advancing toward the gRNA sequence after initial binding, and 2) that gRNA targeting efficiency decreases with a greater amount of 5’ bases transcribed before the gRNA spacer sequence. The first hypothesis on steric hindrance is grounded in the fact that prior works have taken advantage of this strategy to create transcriptional repressors in pol II systems [27] [28] [29]. The second hypothesis on 5’ base interactions can be attributed to potential secondary structure interactions within the transcribed RNA such as hairpin formation, which could interfere with the proper folding of the gRNA scaffold or prevent gRNA recognition of its target sequence.

**Figure 3:**
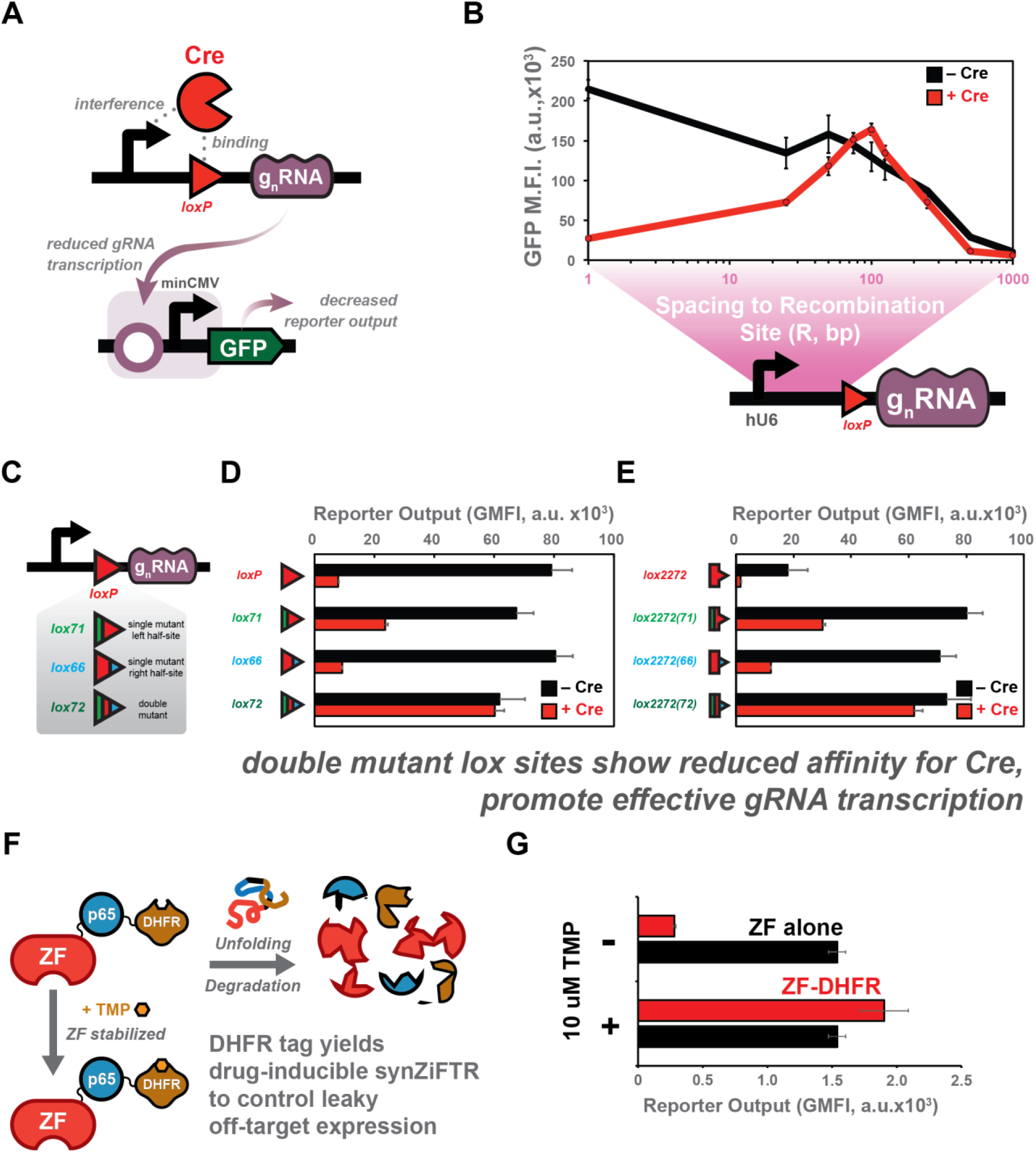
gRNA and synZiFTR expression optimized for use in BLADE decoder circuit template. **A**) SSR binding to target sites post-recombination may inhibit polymerase binding and transcription. **B**) varying the spacing between the transcription start site and SSR target site shows the interplay between 5’ UTR length, SSR inhibition of polymerase binding, and gRNA transcription level. Data represent the fluorescence output signal of a gRNA reporter transfected with different gRNA expressing plasmids containing variable spacer lengths between the TSS and SSR target site/gRNA. **C**) Mutant lox sites of varying affinity for maximizing gRNA production by reducing steric hindrance of SSR binding and minimizing the 5’UTR length. Double mutant lox72 (**D**) and lox2272(72) (**E**) placed between the hU6 promoter and gRNA transcript show similar levels of gRNA reporter output (and decreased affinity for Cre) regardless of SSR addition with no spacer sequence needed between the TSS and SSR target. **F**) synZiFTR off-target expression can be controlled by tagging the TFs with a DHFR domain, stabilized by the addition of TMP in the desired circuit states. **G**) Circuits expressing DHFR-tagged synZiFTRs display significantly less basal expression of TFs in the Z10 and Z10 in the Z11 address. All data represent the geometric mean and standard deviation of flow cytometry data collected from three technical replicates (*n = 3*) of transiently transfected HEK293FT cells collected 48 post transfection.

To address both potential issues, we screened two single mutant target sites (lox66 and lox71) that are reported to maintain a high affinity for Cre (**Figure 3C**) and a double mutant site (lox72) that shows a weak Cre affinity [30] [31] (**Table S2**). Incorporating the double mutant lox72 site between the hU6 promoter and gRNA sequence should 1) lessen the impact of bound Cre preventing polymerase transcription and 2) allow for minimized spacing requirements between the TSS and gRNA sequence. Indeed, the lox72 construct shows comparable gRNA reporter output regardless of Cre expression and with no spacing needed between the TSS and gRNA sequence, while WT, lox66, and lox71 sites show on average a 10x drop in output from a gRNA reporter in the presence of Cre (**Figure 3D**). The same trend is observed by mutating the half-sites of lox2272 to create lox2272(66), lox2272(71), and lox2272(72) (**Figure 3E**). Thus, to efficiently drive gRNA expression from the Layer 1 circuit, the lox2272 and loxP sites are exchanged for lox2272(66)/lox2272(71) and lox66/lox71. When recombined, these single mutant sites become the double lox2272(72) and lox72, allowing for efficient recombination while minimizing SSR binding post-recombination.

We find that the Flp/frt system does not show the same transcriptionally repressive effects as Cre/lox, possibly due to a lower affinity of Flp for its WT frt target site compared with Cre/lox (Data not shown) [32] [33]. As such, we maintain the WT frt sites in the gRNA Layer 1 design. Additionally, we did not mutate Cre or Flp target sites in the synZiFTR system and found that there is minimal change in reporter output between constitutively expressed synZiFTRs with no SSR target sites and synZiFTRs expressed from a recombined BLADE circuit containing WT SSR target sites (**Figure S2a**). This is likely due to the need for translation of these TFs prior to targeting their promoter, thus decreasing the impact of SSR target sites in the 5’ region of synZiFR transcripts. We do note, however, that gRNA expression from any plasmid containing SSR target sites between the TSS and gRNA spacer region diminishes reporter output from cognate gRNA promoters (**Figure S3a**). We attempted to resolve this issue by placing a cleavage site for the RNA processing enzyme CasE between the SSR target sites and the gRNA spacer sequence, but observe little difference in gRNA-promoted reporter output (**Figure S3b**). While this observation does not functionally inhibit the gRNA system, it does limit the amount of overall expression that can be obtained from it.

When assembling the synZiFTR Layer 1 circuit we observe significant leaky expression of TFs in both the Z_01_ and Z_10_ addresses when selecting the Z_11_ address. This leaky expression is likely due to transient expression of Z_01_ and Z_10_ as the Layer 1 circuit is cleaved by either Cre or Flp before being cleaved by the second SSR to reveal Z_11_. To alleviate this issue, we screened circuits containing synZiFTRs in addresses Z_01_ and Z_10_ tagged with either degron (PEST [34] and IKZF3 [35] [36]) or destabilization (dihydrofolate reductase (DHFR) [37]) domains to minimize their expression during these transient stages of incomplete circuit cleavage. PEST and IKZF3 domains promote degradation via the ubiquitin pathway, and IKZF3-mediated ubiquitination is induced by binding to the small molecule drug lenalidomide. The DHFR domain promotes protein unfolding and degradation, and is stabilized through binding to the antibiotic trimethoprim (TMP) (**Figure 3F**). Both PEST and IKZF3 tags helped to reduce transient protein expression for one but not both intermediate addresses (**Figure S2b-e**), whereas DHFR-tagged synZiFTRs display near complete silencing (**Figure 3G**) without the addition of TMP and high induced expression in the presence of TMP. We hypothesize that the DHFR domain performs better than the degron-based systems due to its destabilizing nature to promote rapid protein degradation, rather than relying on protein ubiquitination alone to initiate degradation.

### Libraries of inducible promoters enable tailored circuit output levels

A significant challenge this work seeks to address is the inability to couple robust logic computation with tunable levels of circuit output. Many circuits benefit from the digitization of an analog input signal [21] [22] [23] [24], as it allows for the propagation of signals through multiple components with high signal and low noise. However, natural biological signals operate in the analog space, where different levels or combinations of input signals yield variable levels of the output response. It is thus critical to provide a method to convert digital signals into analog responses. Traditional SSR-mediated transcriptional logic gates yield a high dynamic range between on and off states but lack the ability to finely regulate the level at which that output is produced. To improve upon previous multi-layer circuit designs and translate them to mammalian systems [18] [38], CREATE connects circuit inputs (SSRs) and outputs (fluorescent reporters) via transcription factor wires rather than direct SSR-mediated excision and inversion logic gates in human cells. This allows us to convert digital SSR input signals into analog outputs by defining how strongly each transcription factor activates a cognate promoter.

To ensure our devices produce a wide range of output strengths, we designed, built, and characterized a library of promoters for each programmable gRNA and synZiFTR. gRNA promoters are designed by placing operator sites complementary to each gRNA spacer sequence on the 5’ end of a minimal CMV (minCMV) core promoter. We explored a range of parameters hypothesized to affect promoter strength, including the 1) number of gRNA operator sites, 2) spacing between operator sites, and 3) distance between operators and the minCMV promoter. We find that increasing the number of operator sites upstream of the core promoter yields increasing transcriptional output up to 6x operators, with diminishing transcriptional returns with increased sites (**Figure S4a,b**). Placing the operator sites compactly yielded relatively weak transcriptional output (**Figure S4c,d**), and we note that >6 bp intra-operator spacing is necessary to maximize the reporter signal. This effect is likely due to steric hindrance associated with multiple bound TFs in close proximity. We also note that placing the operator sites further from the core promoter leads to a consistent decrease in promoter strength (**Figure S4e,f**). After elucidating the design rules for gRNA-responsive promoters, we fabricated a suite of promoters for each gRNA capable of driving a wide range of transcriptional output strengths through single-base mutations at each position of a single gRNA operator upstream of the minCMV promoter (**Figure S5**)(**Table S3**). Screening these mutant operator sites alongside promoters containing 1x, 2x, and 3x non-mutant operator sites shows a dynamic range of up to 8.5-fold, 6-fold, and 44-fold between the strongest and weakest g1, g8, and g13 promoters, respectively (**Figure S6a**).

The synZiFTR promoter library was fabricated based on previously defined design rules [26], consisting of zinc-finger operator sites upstream of a minimal TATA box core promoter [39]. We built promoters containing up to 8x operator sites that display increasing transcriptional output with increasing operator sites, unlike the gRNA promoters which we observe limited expression changes after 6x operators. Each synZiFTR promoter library demonstrates up to 14.5-fold, 16-fold, and 16-fold dynamic range between the strongest and weakest ZF1, ZF5, and ZF6 promoters, respectively (**Figure S6b**). Due to the wide dynamic range achieved by modulating the number of zinc finger operator sites within each promoter, we did not study the effects of mutating these operator sites similar to our approach to generating the gRNA promoter library.

Our library of promoters offers two unique capabilities: 1) rapid generation of new circuits by modulating the output and number of TF-responsive promoters in a single cell and 2) grouping series of promoters responsive to each TF that predictably vary the expression of the same output protein from a single cell (**Figure 4A**). To demonstrate the former, we assembled 189 unique 2-input-3-output circuits expressing combinations of GFP, tdTomato, and BFP outputs. These circuits are constructed by co-transfected a Layer 1 TF circuit with unique combinations of promoters expressing different fluorescent proteins that respond to each TF. Each TF from the Layer 1 circuit can target up to three unique outputs, specified by the truth table in **Figure 4B**. Without optimization, ∼90% of circuits perform as expected based on a vector proximity measure (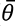 < 15°, [5] [40]).

**Figure 4:**
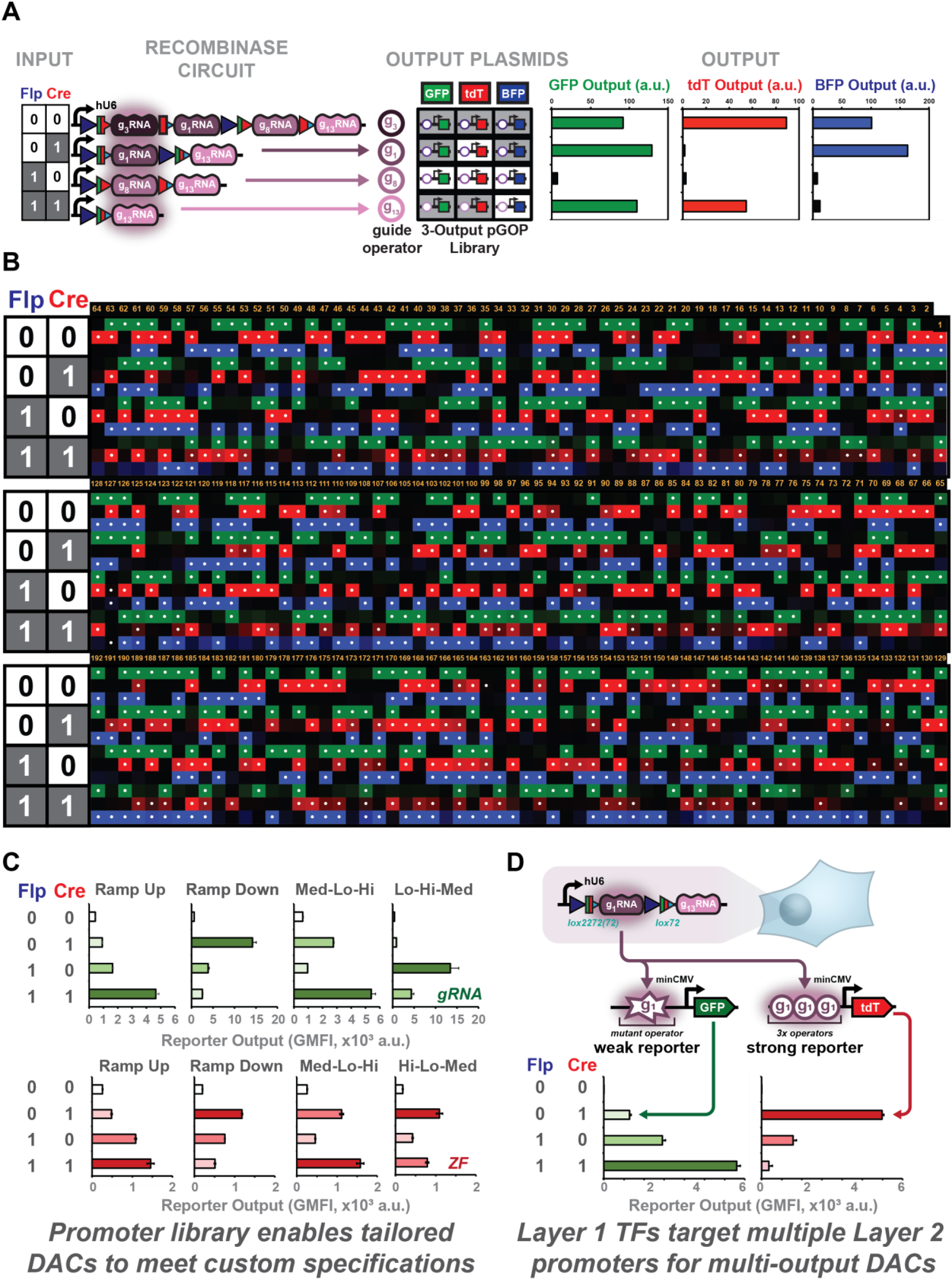
The CREATE platform and a library of promoters enable highly programmable output signals and multiple digital-to-analog circuits in the same cell. **A**) The Layer 1 TF Decoder circuit can be combined with different combinations of Layer 2 promoters expressing different fluorescent outputs, each responsive to a unique TF. Different combinations of Cre and Flp inputs to the circuit specify the TF that is expressed, which then targets the Layer 2 promoter to generate an output signal, with an example flow cytometry data of a 2-input-3-output circuit shown at left. **B**) 192 circuits generated by pairing the Layer 1 circuit with different combinations of Layer 2 outputs expression GFP, BFP, and tdTomoato reporters.189 are unique and 90% function as intended without optimization, quantified using a vector-proximity metric (see supplemental methods). White dots over a square indicate an address that should be active (*i.e.* has an observable signal), and the brighter the coloration the stronger expression observed from the address. **C**) Multiple gRNA (top, green) and synZiFTR (bottom, red) DACs display unique expression patterns and levels of output for each circuit type measured via flow cytometry. Layer 1 circuits are transfected with combinations of characterized Layer 2 promoters intended to yield the desired expression pattern. Combinatorial addition of Cre and Flp SSRs are then added to select for different TF addresses from Layer 1 and the corresponding level of fluorescence output from Layer 2. **D**) Multiple DACs can function in the same cell to create two unique expression patterns by co-transfecting a single Layer 1 design with multiple promoters responsive to each selected TF. All data represent the geometric mean and standard deviation of flow cytometry data collected from three technical replicates (*n = 3*) of transiently transfected HEK293FT cells 48 hours post transfection.

It has been difficult to implement digital-to-analog systems in mammalian cells, which transduce Boolean input signals (SSRs) into a graded output signal, due to a lack of well-characterized constitutive promoters with a sufficiently wide range of transcriptional output. The CREATE platform is uniquely positioned to address this problem through its use of programmable TFs as molecular wires coupled with an extensive library of promoters offering a wide range of output expression strengths. After our initial experiments characterizing the output levels from the promoter library coupled with gRNA or synZiFTR Layer 1 circuits, we selected Layer 2 promoters responsive to each address (TF) of the Layer 1 circuit in combinations hypothesized to yield different patterns of DAC reporter expression. We transfected these groups of Layer 2 promoters with the Layer 1 TF circuit and all possible combinations of Cre and Flp recombinases to select for different levels of reporter output into HEK293FT cells. For both gRNA and synZiFTR systems, the CREATE platform demonstrates reproducible digital-to-analog circuits with a range of behavior, such as ramp up (increasing output) and ramp down (decreasing output) trends (**Figure 4C**). These behaviors are programmed simply by selecting promoters of a desired transcriptional strength for each circuit address. It is also possible to regulate unique patterns of expression for multiple proteins simultaneously by transfecting two sets of DACs in the same cell, a task difficult to program for SSR-based systems alone. To do so, each Layer 1 TF targets two separate promoters in Layer 2 yielding the desired expression profile of each unique output. We demonstrate this by ramping up GFP reporter levels while simultaneously ramping down tdTomato reporter levels in HEK293FT cells (**Figure 4D**).

### Executing complex operations by wiring circuits in parallel

In addition to single outputs, CREATE can accommodate sets of 2-input-4-output circuits in Layer 2. This can be thought of as a system of parallel processors, where a complex operation is distributed such that smaller computations by individual processors contribute to the function of a larger system. To demonstrate the computational power of this approach, we develop a genetic multiplier circuit in a human cell line (HEK293FT). This device computes the product of two 2-bit inputs (combinations of SSRs) to yield a unique combination of fluorescent reporter outputs (**Figure 5A**). Transcription factors selected by Cre and Flp inputs in Layer 1 target their cognate promoters in Layer 2, which yield unique outputs selected by combinations of VCre and KD inputs, as demonstrated in **Figure 5B**.

**Figure 5:**
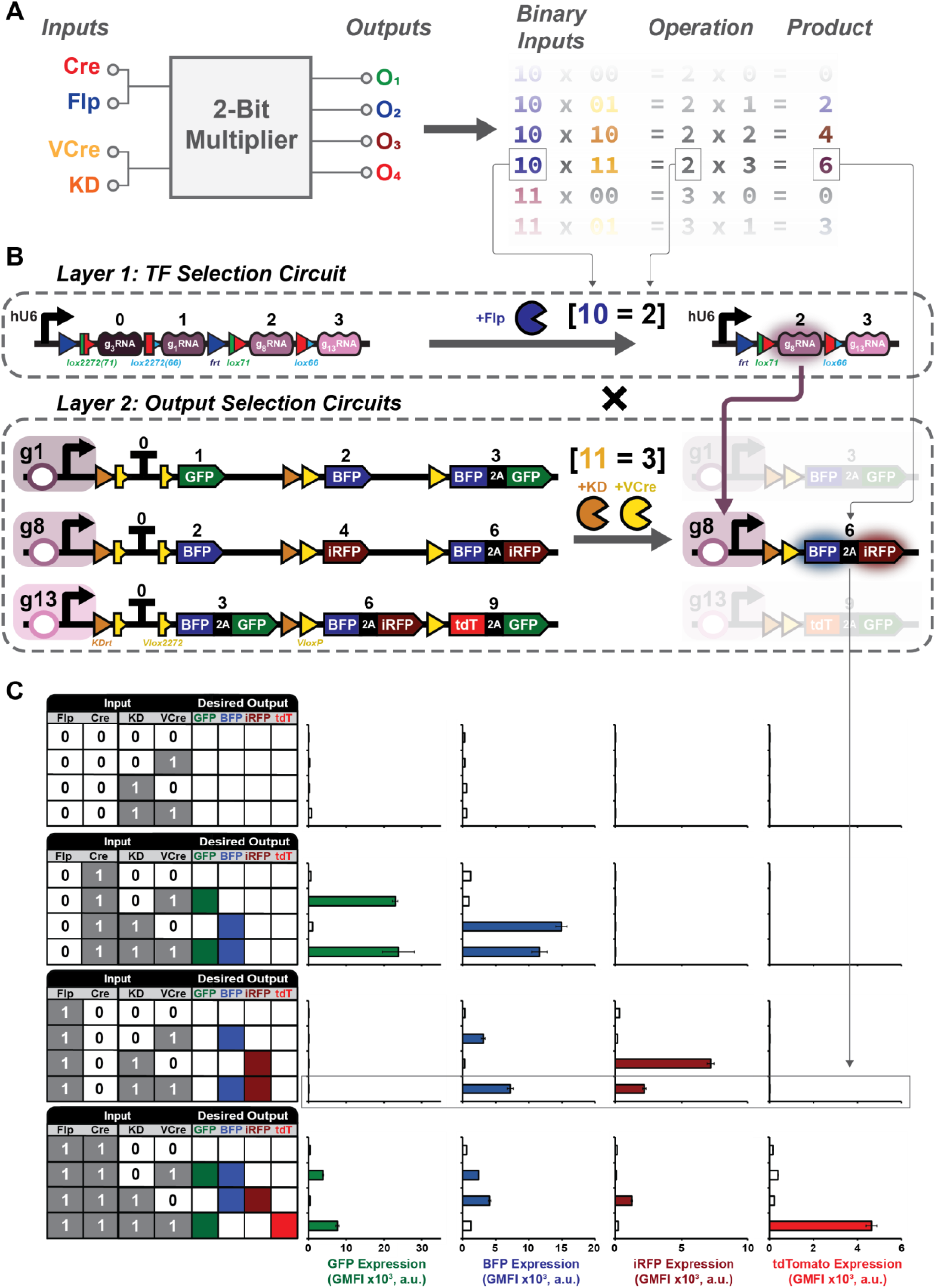
Connecting multiple circuit layers via TF wires enables 4-input-16-output circuits with advanced computational capabilities. **A**) Abstract representation of the genetic multiplier circuit. Our circuit takes two 2-bit recombinase inputs (Cre | Flp, and VCre | KD) and computes the product of these inputs to yield a unique fluorescent output. On the right, example calculations are shown where the binary input signals are processed by the circuit to yield the desired output signal. **B**) Biological representation of the genetic multiplier circuit. The multiplier is broken into two separate layers of 2-input-4-output decoder circuits where combinations of Cre and Flp SSRs select a TF from Layer 1 that activates a desired Layer 2 circuit, which is cleaved by VCre and KD inputs. In this example, Flp [binary value of 10 = 2] selects for the expression of g8, and the Layer 2 circuits are cleaved by both VCre and KD [binary value of 11 = 3]. g8 then activates its cognate Layer 2 circuit to compute the operation 2 x 3 = 6, represented by an output of dual BFP and iRFP720 expression. **C**) Flow cytometry data of the distributed multiplier circuit. Input and desired output values for each multiplier address are specified by the truth table at the left, and flow cytometry data is displayed at the right for all fluorescent reporters used. Layer 1 circuits were transfected with individual Layer 2 circuits into separate populations of HEK293FT cells and data represent the fluorescence values from the on-target Layer 2 circuits (see supplement for off-target data). All data represent the geometric mean and standard deviation of three technical replicates (*n = 3*) collected 48 hours post transfection.

All SSRs used in the multi-layer circuit design are observed to be orthogonal to one another [5] (**Figure S7**). Similar to our initial TF orthogonality screen, the multiplier circuit design also shows no cross-reactivity between selected TFs and TF promoters (**Figures S8**) although some leaky expression of gRNAs in addresses Z10 and Z01 is observed when selecting for address Z11. To demonstrate the operation of the multiplier, we transfected the Layer 1 TF circuits with each Layer 2 reporter circuit separately into populations of HEK293FT cells and collected flow cytometry data 48 hours post-transfection. There is a clear distinction between the intended ON and OFF states for each fluorescent reporter (**Figure 5C**), and each state produces the intended operation measured by a vector proximity metric (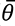 = 2.94°). We note that when all Layer 2 circuits are transfected into the same cell, the basal level of several reporter states was difficult to distinguish from the intended on-target expression levels of the same reporters (**Figure S9**). However, the function of this circuit is quantitatively accurate (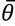 = 2.93°), and would likely perform as intended if there was no need to reuse fluorescent reporters in different states. Both distributed and combined versions of the CREATE platform should be advantageous for many applications, such as drug screening platforms where SSRs are driven from inducible promoters to digitize and amplify effects on biological signals related to the tested compounds [24] [23].

To further stress-test the capabilities of the CREATE platform, we engineer a series of mixed-signal analog and digital state machines in HEK293FT cells by combining the DAC architecture with a 2-input-4-output Layer 2 circuit (**Figure 6A**). Each of the circuits presented here was fabricated simply by combining previously characterized DAC architectures with a 2-input-4-output Layer 2 circuit. Importantly, each circuit generated using this plug-and-play approach demonstrates the intended response profile without the need for iterative optimization. Cells were transiently transfected with a ramp-up GFP DAC and gRNA 8 responsive circuit along with all combinations of Cre, Flp, VCre, and KD specified by the truth table in **Figure S10**. This circuit demonstrates the conversion of digital SSR inputs to analog GFP outputs convolved with digital BFP and iRFP reporter signals. Additionally, we demonstrate the regulation of multiple DAC configurations using a hi-lo-med GFP DAC and a lo-hi-med tdTomato DAC transfected with the 2-input-4-output gRNA 8 responsive circuit (**Figure 6B**). While we note a minor loss of fidelity for the tdTomato low state, all other aspects of the mixed digital and analog signal generator appear intact. The loss of tdTomato reporter expression in the low state may be attributed to resource competition, as the promoter used to drive this output is weak initially. It also draws away transcriptional resources (gRNA/dCas9-VPR complexes) using a second hi promoter, the transcriptional output from the tdTomato reporter may be too weak to generate a sufficient signal. We posit that this problem can be easily addressed by selecting another weak promoter from our gRNA promoter library for use with the tdTomato lo reporter.

**Figure 6:**
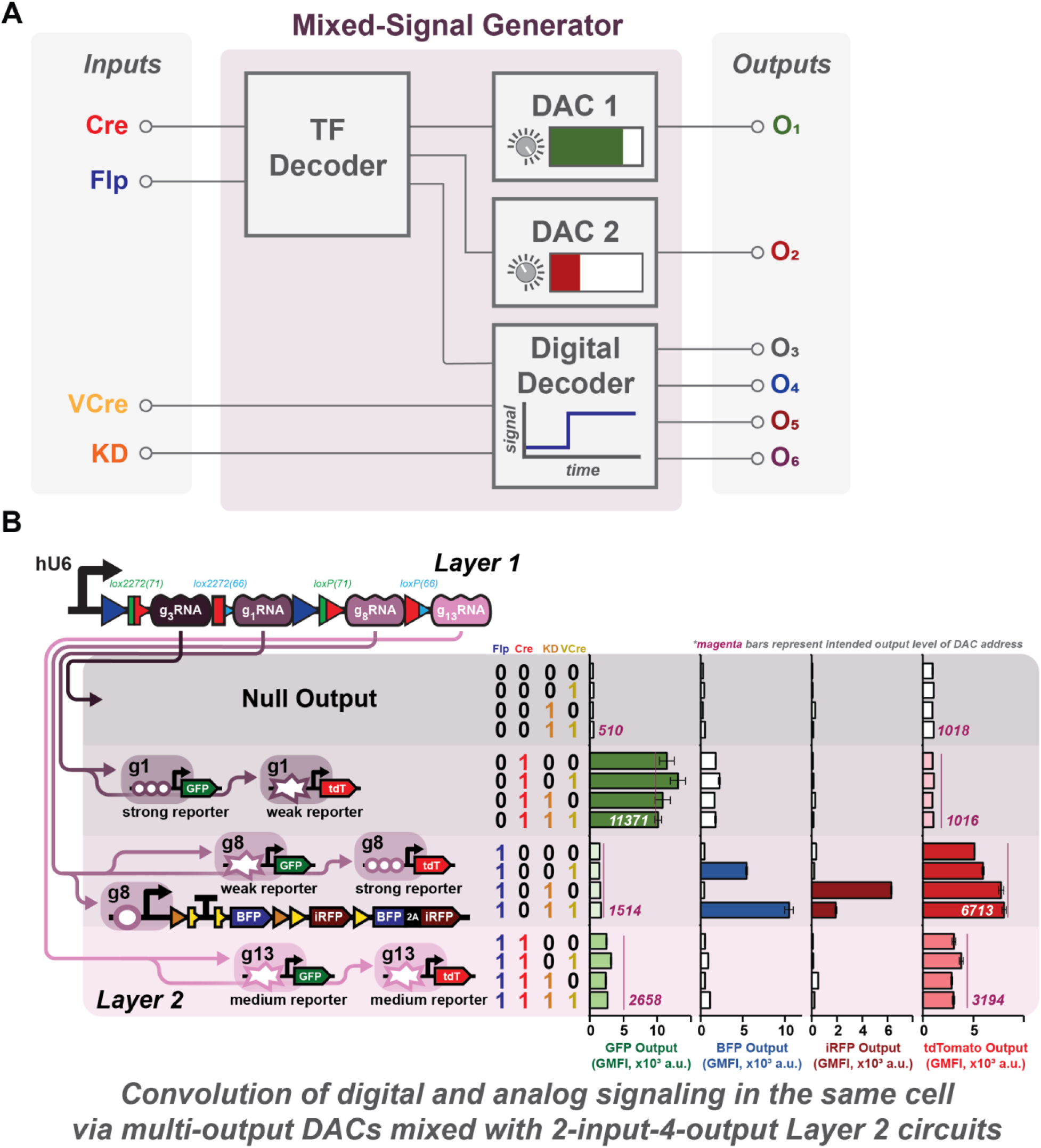
Mixed-signal generation in single cells achieved by combining DACs and Layer 2 output circuits. A) Mixed-signal circuit abstraction. To design a mixed-signal generator capable of producing both digital and analog output signals, the Layer 1 TF decoder is connected to multiple Layer 2 circuits consisting of outputs driven by promoters of varying transcriptional strength (DACs) and 2-input-4-output decoder circuits whose output is selected by combinatorial addition of VCre and KD SSRs. **B**) Wires connecting Layer 1 TFs with Layer 2 reporters detail the fluorescent proteins and relative signal strengths at each address. Address inputs are specified by the truth table immediately to the left of the data and lighter and darker shades of color indicate addresses expected to be relatively weak or strong, respectively. Cells are transiently transfected with all circuit components (DAC 1, DAC 2, Digital Decoder, and TF Decoder) into HEK293FT cells and unique combinations of SSR inputs are added to different transfection groups to select for different circuit states. High-low-medium (GFP) and low-high-medium (tdTomato) DACs each show consistent reporter expression when Layer 2 SSRs are added but variable expression when Layer 1 SSRs selected for different TFs. The addition of a 2-input-4-output g8 responsive BFP/iRFP reporter circuit yields the intended outputs as well and does not appear to negatively impact the function of the DAC circuits. All data represent the geometric mean and standard deviation of flow cytometry data collected from three technical replicates (*n = 3*) 48 hours post transfection.

## Discussion

Genetic circuits provide cells with the instruction to perform complex regulatory tasks. Synthetic gene circuits have thus far focused on the implementation of either digital or analog circuits, but would benefit from a unified and adaptable framework to program both modes of signal processing and propagation. Our multilayer CREATE platform can integrate multiple input signals to yield the desired pattern of convolved digital and analog output signals in mammalian cells. CREATE builds upon existing site-specific recombinase (SSR) logic circuits by incorporating programmable transcription factor (TF) ‘wires’ between circuits in parallel. The system is developed to interface with both CRISPRa and zinc-finger transcription factor systems, where a unique set of SSR inputs to Layer 1 selects the desired TF to initiate transcription in Layer 2. The CREATE platform robustness and modularity of this new technology by building 189 unique 2-input-3-output logic circuits and show that 90% perform their intended operations without optimization. Additionally, we present the first genetic multiplier circuit in mammalian cells, demonstrating the computational power of parallel processing when performing complex operations on multiple input signals. CREATE is also capable of fine control over output gene expression levels, yielding eight different digital-to-analog circuits with no optimization required, the first multi-output DAC circuit in mammalian cells, and the only mixed digital and analog signal generator developed to date.

Synthetic biological signaling in the analog space often suffers from a loss of signal as it is transferred between circuit elements. Previous work has shown that digitizing these signals can lead to amplification of weak input signals [23] [24] [22] [20] [21] and allows for high fidelity information transfer despite passing multiple circuit modules. By coupling digital processing circuits with an analog output signal in this work, we expand the use of digitizer modules to settings where analog input signals are received, and analog output signals are required. An additional benefit of the CREATE platform is its use of recombinase logic to drive such analog outputs, creating a memory device in cells. This type of memory response would be useful for differentiating cells toward a particular phenotype defined by differential expression of the same protein, such as varying FOXP3 expression in naïve and regulatory T cell subsets [41] [42] [43] [44] or adjusting chimeric antigen receptor expression to match different antigen density profiles [45] [46].

CREATE decouples logic computation (SSR inputs) from circuit output (TF-induced transcription). This allows modulation of the output level by altering the TF or TF-targeted promoter while still performing logic computation with the precision and memory afforded by SSR-mediated excision reactions. To maximize control over output expression level, we present a library of gRNA and zinc-finger responsive promoters. Our library exhibits a wide range of transcription strengths to enhance the user-defined performance of circuits developed with the CREATE platform. By coupling these promoters with a Layer 1 circuit, we demonstrate 189 unique and modularly assembled circuits and 4 digital-to-analog circuits for each TF system with no optimization required. Additionally, by combining promoters of different strengths that respond to the same TF, we show the differential regulation of multiple protein expression levels in the same cell.

Wiring together multiple single-layer circuits using transcription factors greatly expands the operations a system may perform. Previously developed single layer SSR circuit designs display robust logic computation but are limited by the number of heterospecific recombination sites available for each SSR [4] [5] [8]; building a single layer multiplier circuit like the one shown in this work (or any 4-input-16-output circuit) would require up to 8 unique SSR target sites in a single layer design. By breaking this design up into individual 2-input-4-output circuits, these constraints are eased, and the limiting factor becomes the number of orthogonal TFs available. By designing each layer of the CREATE platform to interface with programmable gRNAs and synZiFTRs, the requirement for orthogonal TFs is minimized given the proven methods of designing new DNA-binding domains for each system.

When assembling the multiplier circuit, we note that it becomes difficult to separate intended outputs from targeted Layer 2 reporter circuits from the cumulative sum of basal expression from off-target Layer 2 circuits. While VP metric analysis reveals that this circuit performs generally as expected (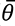 = 2.93), it is visually difficult to separate populations of cells intended to express a reporter in weaker-expressing states from the basal activity of other states. This may be addressed in future iterations through several strategies. Using a stronger transcriptional activator to boost the signal from on-target reporters may increase their signal to a point at which it is clearly distinguishable from any noisy basal expression. Conversely, reducing the level of basal reporter transcription by exchanging the current minCMV and ybTATA core promoters for minimal promoters that display less basal activity would help to address this problem by reducing the amount of off-target transcription. Tagging circuit outputs from Layer 2 with degron tags, such as a PEST sequence, would similarly yield a more rapid turnover of output proteins and may reduce off-target noise. This constraint is specific to the multiplier design, where certain fluorescent protein outputs are reused at multiple addresses. The same design constraints may not be observed for a more general multiplexer circuit, where each output would ber unique.

We envision this platform as a highly versatile tool for modulating complex expression patterns of multiple genes simultaneously, and a way to rapidly program higher-order computational operations in mammalian cells. In this work, we modulate different programs of expression for different fluorescent reporters in the same cell using the CREATE platform. However, these reporter genes can easily be exchanged with other genes of interest, such as cytokines [47] [48] or surface receptors [10, 49] [50] [51], to study how simultaneously regulating the expression level of multiple proteins that affects cell signaling or developmental processes.

The CREATE platform can have many applications, such as bioelectronic interfaces, biological hazard monitoring devices [52], or animal model developments ([53]). It can also be used in theragnostic, especially in the context of irradiated mammalian cells. Irradiating cells prior to infusion has allowed several clinical-stage companies to pursue the translation of immortalized cell lines for cancer therapy [54]. In this strategy, cells are treated with a lethal dose of radiation prior to infusion into patients. The irradiated cells continue to perform their intended biological function for several days in the patient after which time they die off, leaving the patient free of engineered cells but reaping the benefits of their activity. This strategy may enable the translation of the CREATE platform for to generate a highly targeted and poly-functional therapeutic response.

## Acknowledgments

We thank Divya Israni, Huishan Li, and Ahmad Khalil for the kind gift of the synZiFTRs used in this work.

## Author contributions

Project conceptualized by JHL, BHW, and WWW. Plasmid design, cloning, investigation, and data collection were performed by JHL, BHW, MCM, JMM, WKJB, RP, KD, SK, and AP. Manuscript prepared by JHL and edited by JHL, BHW, and WWW.

## Data and materials availability

Data is available upon request. All plasmids developed for use in this work are available through the non-profit plasmid repository Addgene.

## Supplementary Materials

### Materials and Methods

#### Cell lines and culture conditions

Experiments were performed using the HEK293FT cell line purchased from ATCC. HEK293FT cells were subcultured in DMEM medium (Corning) supplemented with 5% heat-inactivated fetal bovine serum (FBS) (Gibco), 50 IU/mL penicillin/streptomycin (Corning), 2 mM L-glutamine (Corning), and 1 mM sodium pyruvate (Lonza) (5PSGN). During cell passaging, 0.05% or 0.25% Trypsin, 0.53 mM EDTA (Corning) was used to detach cells from culture flasks. Trypsin was neutralized using 5PSGN after a 3-minute incubation at 37 °C, and cells were spun down at 300 xg for 5 minutes. After pelleting, medium + trypsin was removed and cells were resuspended in fresh 5PSGN before plating in a new flask.

#### Cell plating, plasmid transfection, and drug induction

24 hours prior to transfection, HEK293FT cells were trypsinized and seeded on 48w plates at a density of 75k cells/well in 250 μL 5PSGN medium. DNA mixes were prepared the day of transfection. 1000 ng of DNA (20 μL DNA at 50 ng/μL) was mixed with 30 μL 0.15 M NaCl solution, and subsequently mixed with 50 μL of a PEI in 0.15 M NaCl. Linear PEI (Polysciences 23966, MW = 25k) was dissolved with the assistance of hydrochloric acid and sodium hydroxide to a concentration of 0.323 g/L in deionized water, filter sterilized (0.22 um), and stored at -80 °C. Stocks were thawed and mixed with 0.15 M NaCl in an 8:42 PEI:NaCl ratio for each transfection mix. Transfection mixes were incubated 15 minutes at room temperature after which 25 μL was added to each of three wells (*n = 3*). DNA components for all DAC and multiplexer circuits tested can be found in **Tables S4-14**.

Trimethoprim (TMP) (Thermo Scientific, J6305303) and Lenalidomide (LLM) (Tocris Biosciences, 6305/100) were each dissolved in dimethylsulfoxide (DMSO) to 1000x stock concentrations of 1 mM and 10 mM, respectively. Trimethoprim stocks were maintained at 4 °C, and lenalidomide stocks were stored at -80 °C. For samples requiring drug induction, 56x stocks were made by mixing the 1000x drug stocks in 5PSGN. From the 56x stocks, 5 μL of the appropriate drug was added to each well for a final working concentration of 1x (1 μM TMP, 10 μM LLM) in 280 μL total volume in each well of a 48w plate.

#### Flow cytometry data collection and analysis

All flow cytometry data was collected using a four-laser Attune NxT Flow Cytometer (Life Technologies) and attached Autosampler. Fluorescence data was collected using the following configuration: eGFP (488 nm excitation laser, 530/30 emission filter), mtagBFP2 (405 nm excitation laser, 450/50 nm emission filter), tdTomato (561 nm excitation laser, 585/16 emission filter), iRFP-720 (605 nm excitation laser, 720/30 emission filter), and LSS-mOrange (405 nm excitation laser, 610/20 emission filter). Prior to flow cytometry, cells were trypsinized (50 μL/well), neutralized in 5PSGN (75 μL/well), transferred to 96w U-bottom plates and mixed thoroughly by pipetting.

Data analysis was conducted using FlowJo software (BD). For each experiment, live cell populations were gated for using an FSC-A vs SSC-A plot and single cells were subsequently gated for on an FSC-A vs FSC-H plot. Transfected cells expressing the LSS-mOrange or iRFP-720 transfection markers were gated for by applying a histogram gate to the top 0.1% expressing cells in a blank, negative control sample pre-processed to display only viable, single cells. Single-positive control samples for each fluorescent protein in the five-color experiments and used to perform compensation.

#### Digital-to-analog circuit design

Each promoter in our custom library was screened for transcriptional activity through transfection with the corresponding Layer 1 circuit and appropriate recombinase inputs to yield their cognate transcription factors in HEK293FT cells. Promoters for each transcription factor system were characterized in the same experiment. The geometric mean of three technical replicates (*n = 3*) was plotted in Excel to determine the relative activity of each promoter under the same inducing conditions. Combinations of promoters responsive to each individual transcription factor were selected based on these screens and were transfected into HEK293FT cells along with all possible combinations of Cre and Flp inputs. A detailed list of promoter combinations corresponding to the DAC data plotted in Figure 4 can be found in **Table S4-11**.

#### *2-input-4-output* circuit design and fabrication

Circuits were assembled using the Unique Nucleotide System (UNS) [55] for Gibson assembly. Each circuit address is first cloned into a ‘part’ vector, containing a constitutive CAG promoter and rabbit beta globin transcription terminator (RBGpA) sequence flanking specific UNS homology regions. Each template part vector (BW701-704, Addgene #87561-87564) is digested with MluI-HF and NotI-HF (New England Biolabs) and the variable insert region is cloned in via Gibson assembly or ligation and verified via test digest using AscI and NheI-HF (New England Biolabs) and Sangar sequencing (Quintara Biosciences). Verified parts are then amplified via PCR using primers listed in **Table S15** and cloned via Gibson assembly into a U1-UX destination vector (BW700, Addgene # 87578) digested with NotI-HF and EcoRI-HF (New England Biolabs). All circuit plasmids used in this work are available from the non-profit plasmid repository Addgene.

#### Vector proximity analysis

A previously established and refined vector proximity metric was used to determine the performance of all digital circuit operations performed in this work [5], [40]. Briefly, circuit data was broken down into two vectors ***t*** and ***s***, where ***t*** represents an ***n***-dimensional binary vector corresponding to the intended truth table of an individual circuit and ***s*** represents the corresponding signal vector composed of fluorescence output values for each address. Recent work by *Bowyer et al.* has shown that the original vector proximity measure can be improved by dividing the original vector proximity metric by the number of fluorescence-expressing addresses (***n_f_***) [40], as it is more difficult for transfected cells to achieve a full population response for fluorescent states than it is for all cells to remain inactive in non-fluorescing states. This modified cosine similarity metric between vectors ***t*** and ***s*** is calculated as

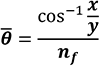

where

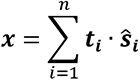

and

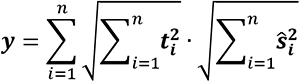

---

**Table S1:**
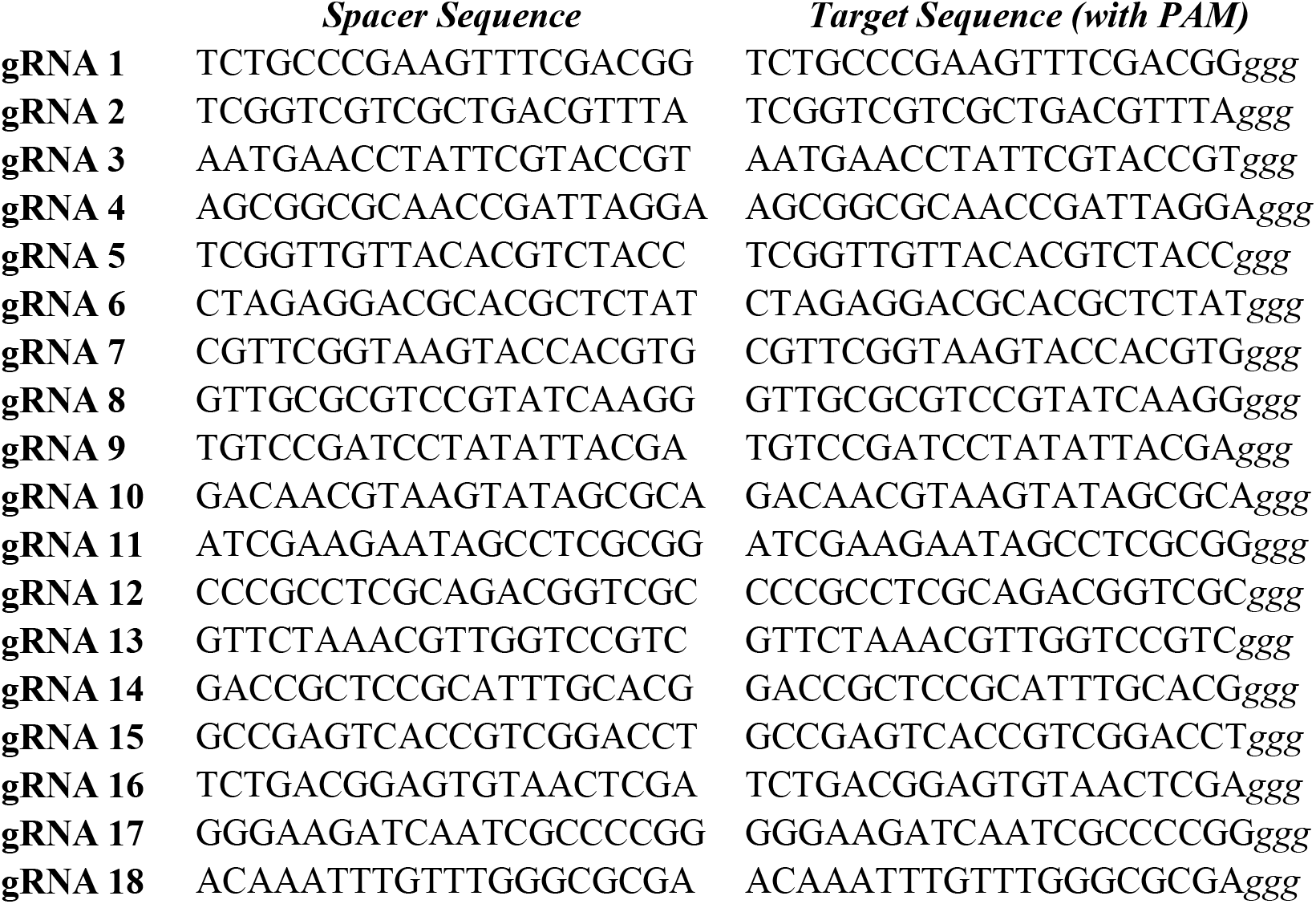
gRNA Spacer & Target Sequences.

**Table S2:**
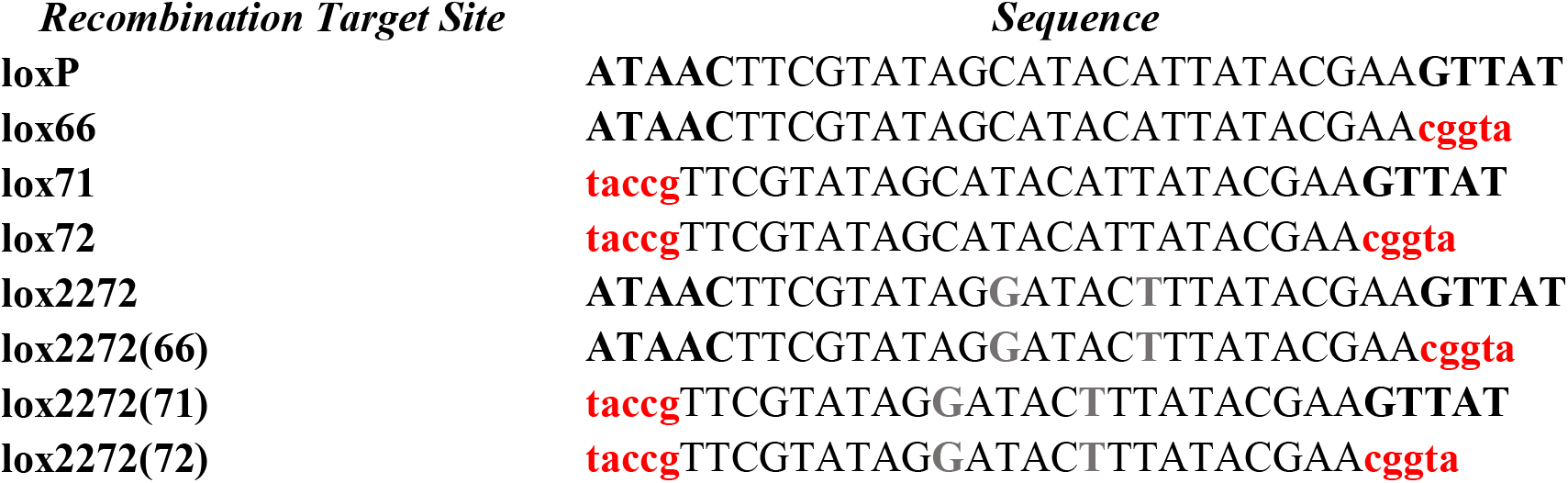

**Table S3:**
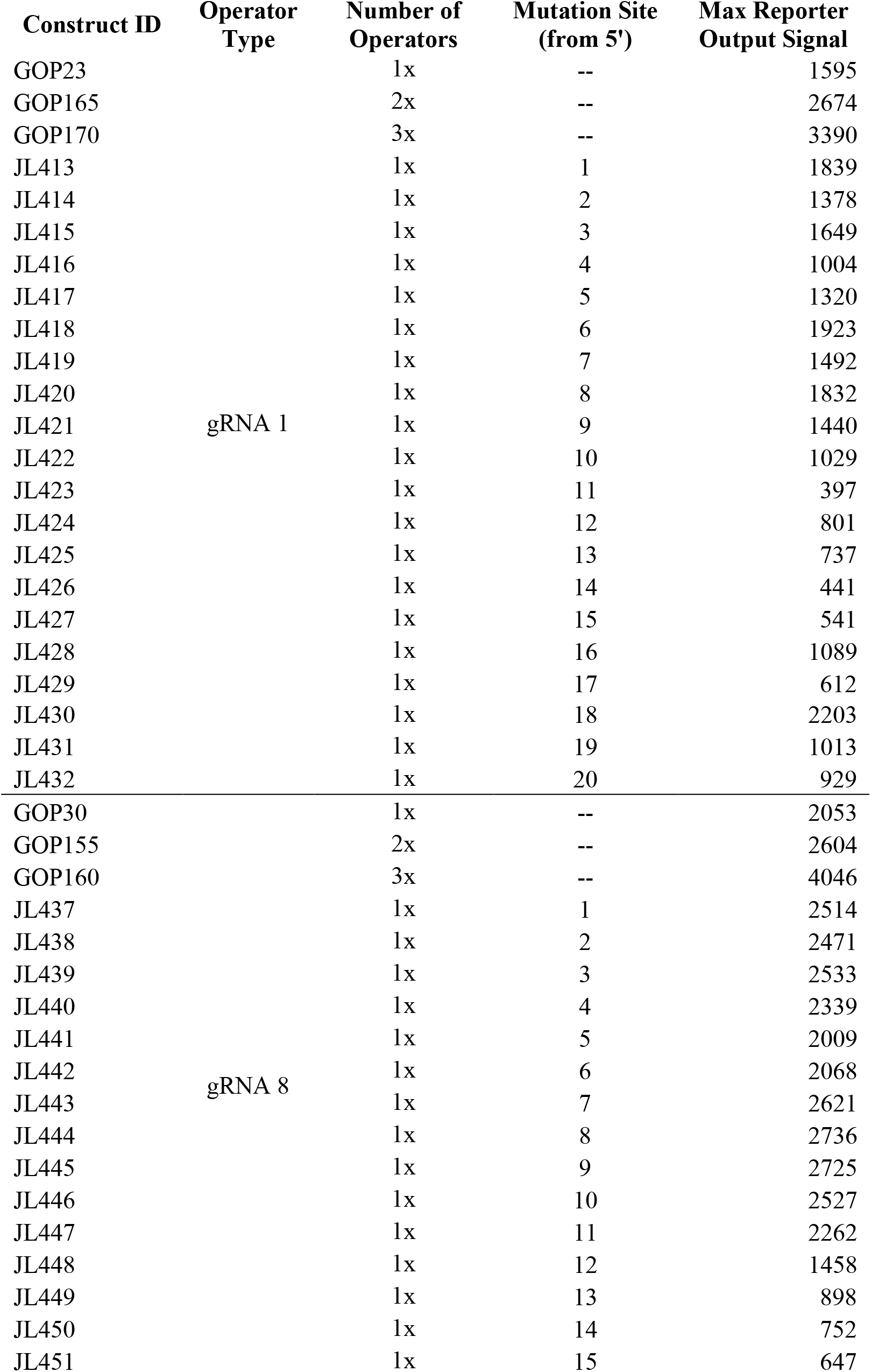
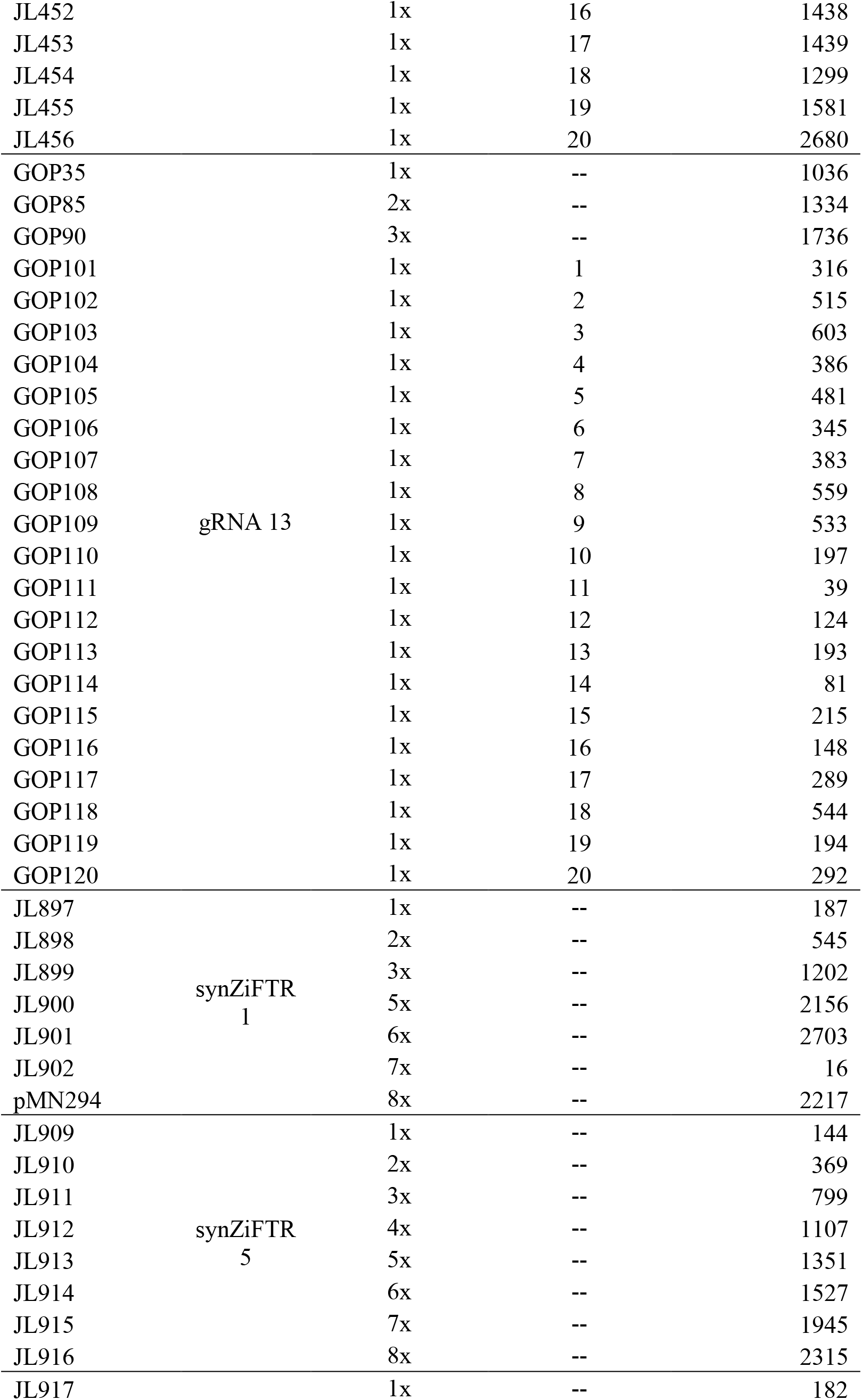
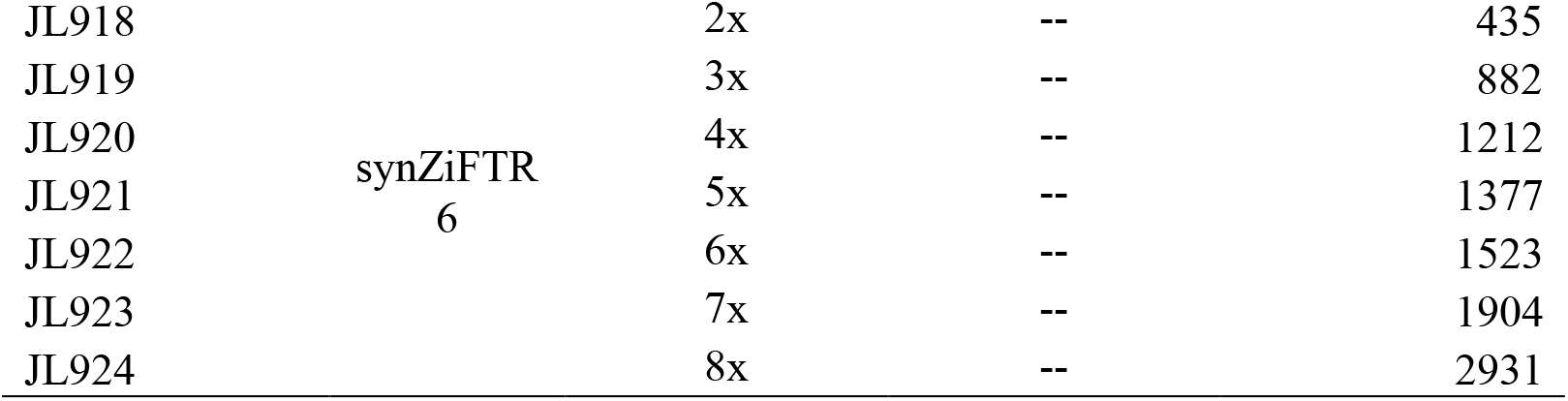
Library of gRNA & synZiFTR Promoters and Relative Transcription Strengths.

**Table S4:**
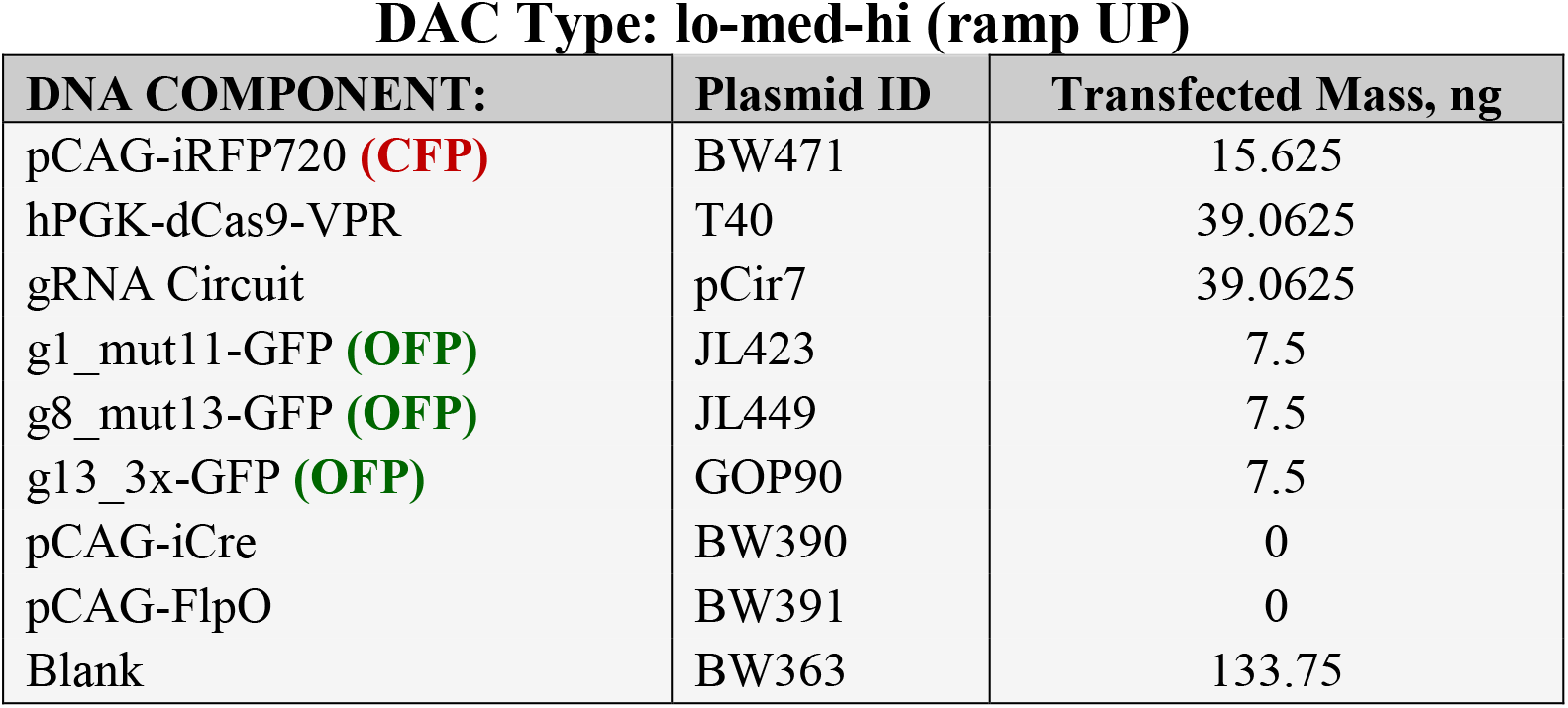
DAC Type: lo-med-hi (ramp UP)

| DNA COMPONENT: | Plasmid ID | Transfected Mass, ng |
| --- | --- | --- |
| pCAG-iRFP720 (**CFP**) | BW471 | 15.625 |
| hPGK-dCas9-VPR | T40 | 39.0625 |
| gRNA Circuit | pCir7 | 39.0625 |
| g1_mut11-GFP (**OFP**) | JL423 | 7.5 |
| g8_mut13-GFP (**OFP**) | JL449 | 7.5 |
| g13_3x-GFP (**OFP**) | GOP90 | 7.5 |
| pCAG-iCre | BW390 | 0 |
| pCAG-FlpO | BW391 | 0 |
| Blank | BW363 | 133.75 |

**Table S5:** DAC Type: hi-med-lo (ramp DOWN)

| DNA COMPONENT: | Plasmid ID | Transfected Mass, ng |
| --- | --- | --- |
| pCAG-iRFP720 (**CFP**) | BW471 | 15.625 |
| hPGK-dCas9-VPR | T40 | 39.0625 |
| gRNA Circuit | pCir7 | 39.0625 |
| g1_mut11-GFP (**OFP**) | GOP170 | 7.5 |
| g8_mut13-GFP (**OFP**) | JL455 | 7.5 |
| g13_3x-GFP (**OFP**) | GOP101 | 7.5 |
| pCAG-iCre | BW390 | 0 |
| pCAG-FlpO | BW391 | 0 |
| Blank | BW363 | 133.75 |

**Table S6:** DAC Type: lo-hi-med

| **DNA COMPONENT:** | **Plasmid ID** | **Transfected Mass, ng** |
| --- | --- | --- |
| pCAG-iRFP720 (**CFP**) | BW471 | 15.625 |
| hPGK-dCas9-VPR | T40 | 39.0625 |
| gRNA Circuit | pCir7 | 39.0625 |
| g1_mut11-GFP (**OFP**) | JL423 | 7.5 |
| g8_mut13-GFP (**OFP**) | GOP160 | 7.5 |
| g13_3x-GFP (**OFP**) | GOP90 | 7.5 |
| pCAG-iCre | BW390 | 0 |
| pCAG-FlpO | BW391 | 0 |
| Blank | BW363 | 133.75 |

**Table S7:**
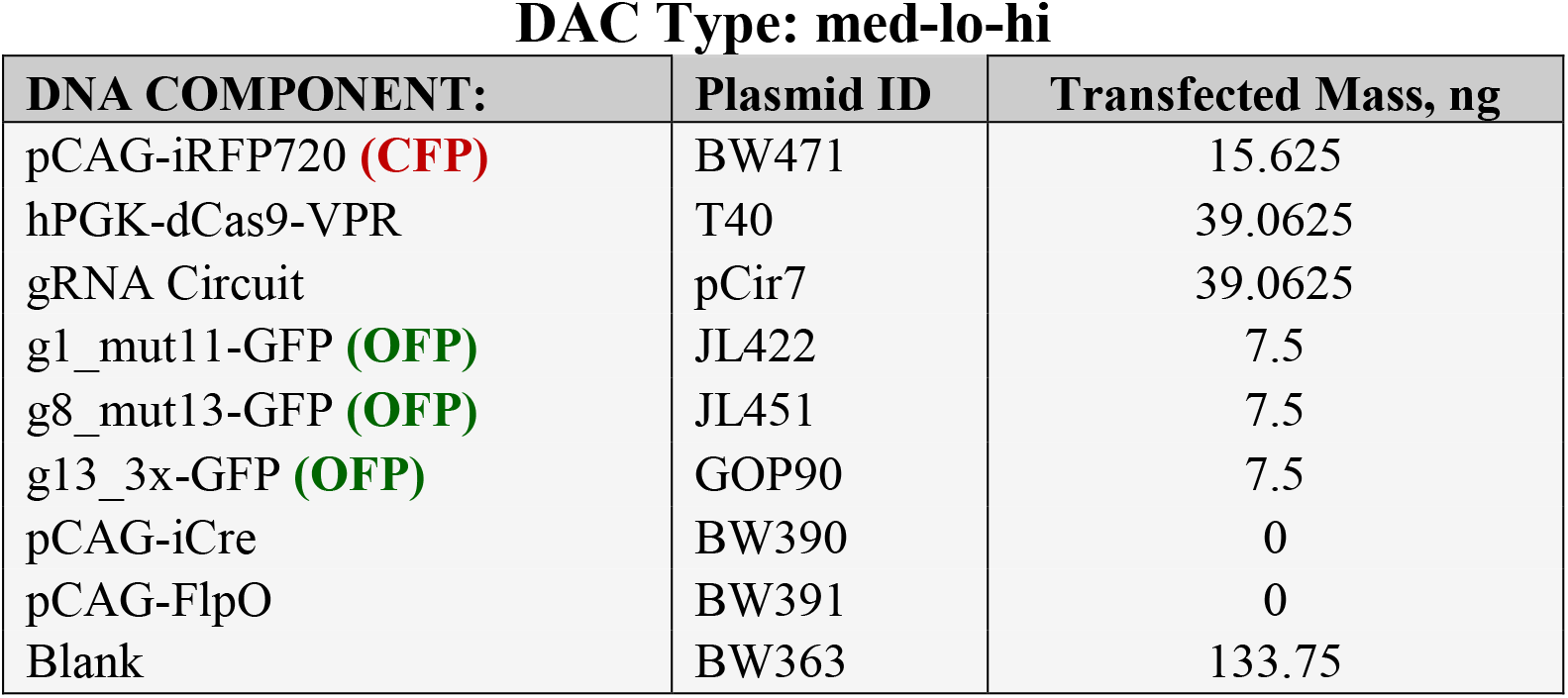

**Table S8:**
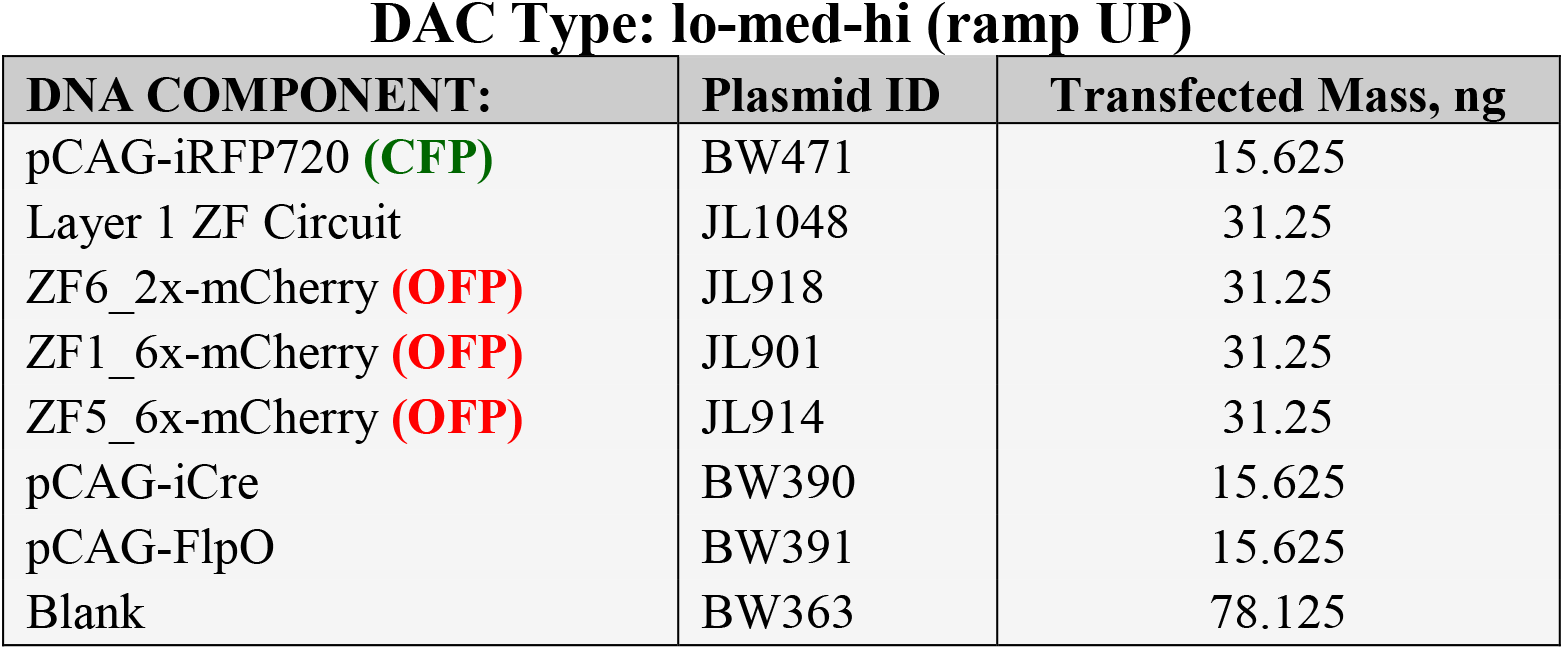

**Table S9:**
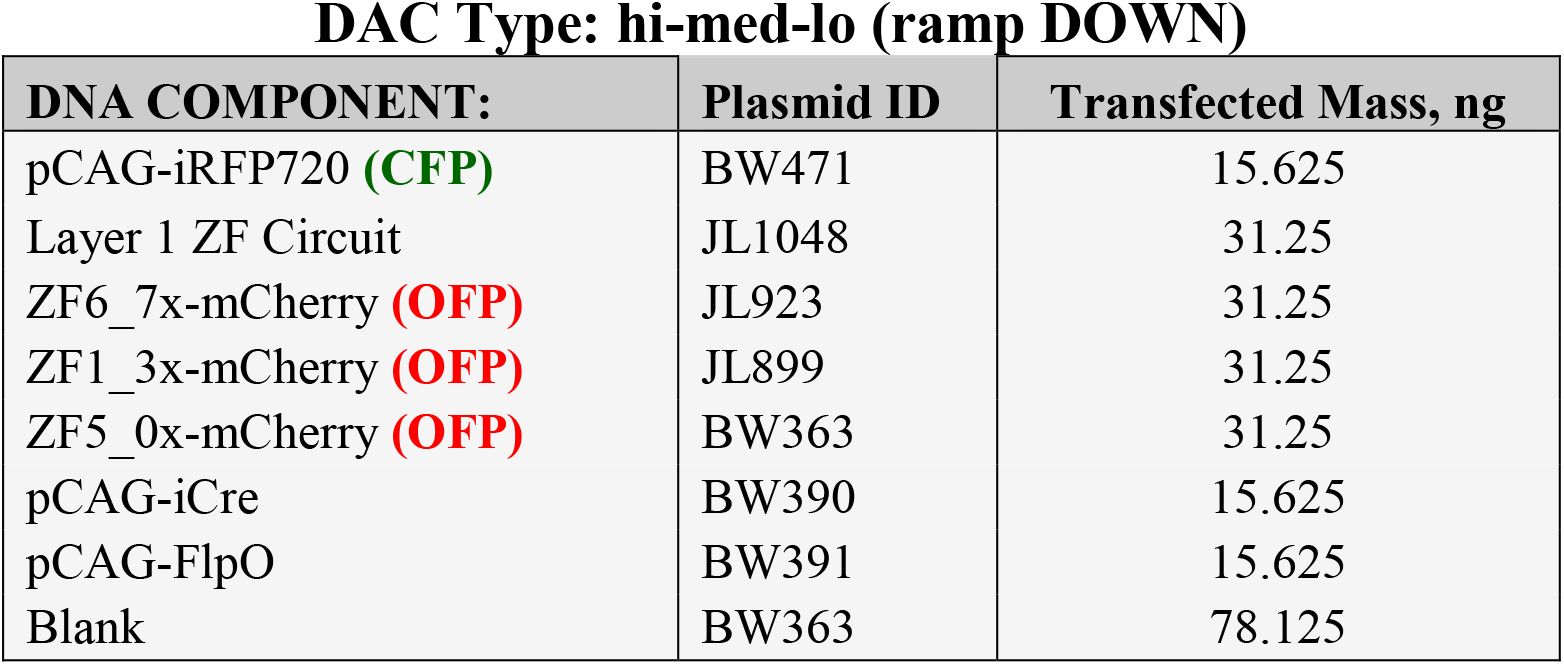

**Table S10:**
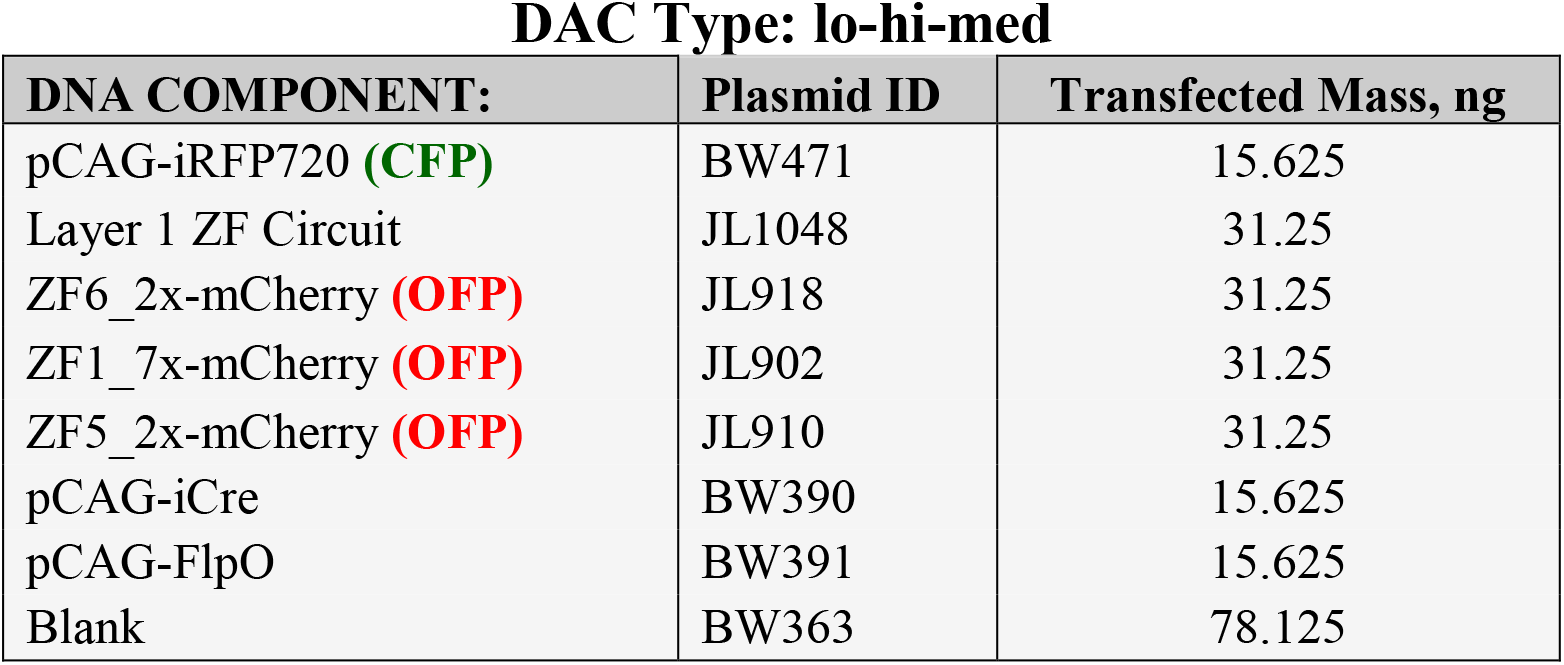

**Table S11:**
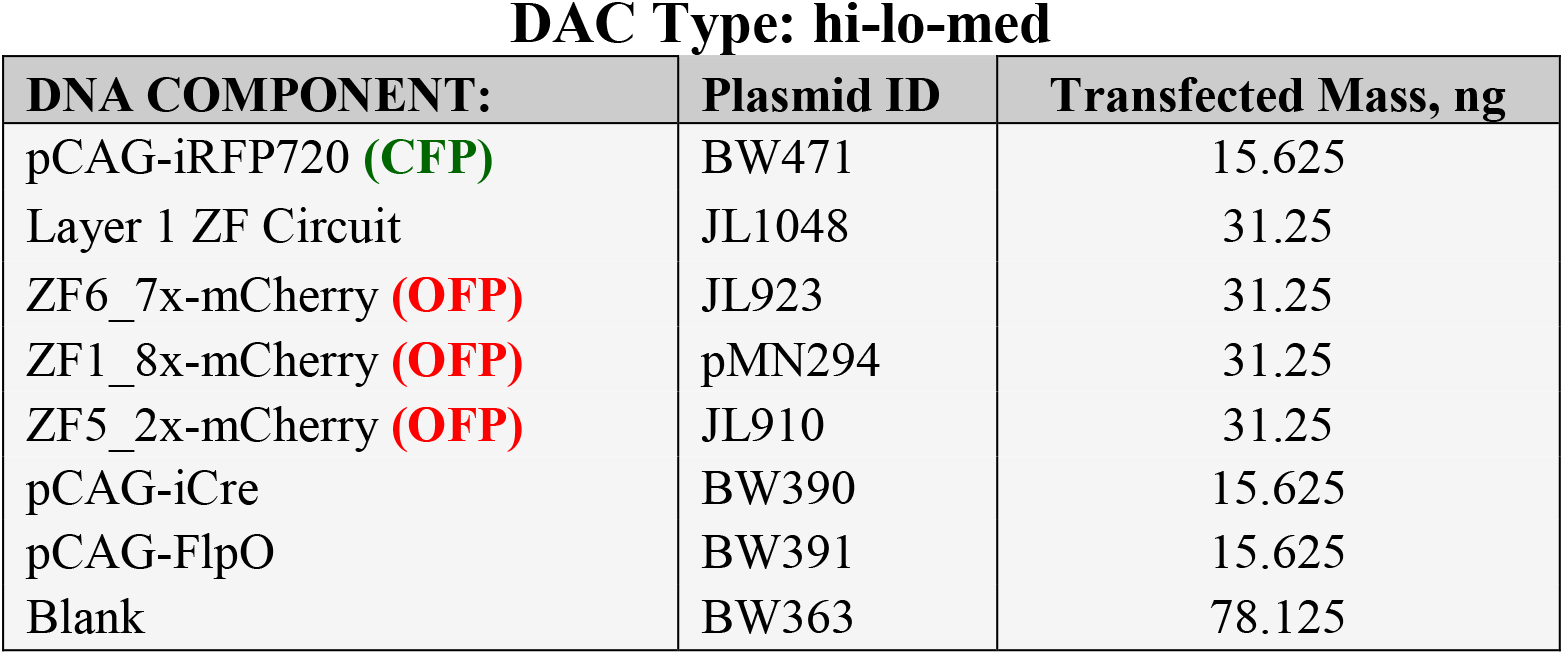

**Table S12:**
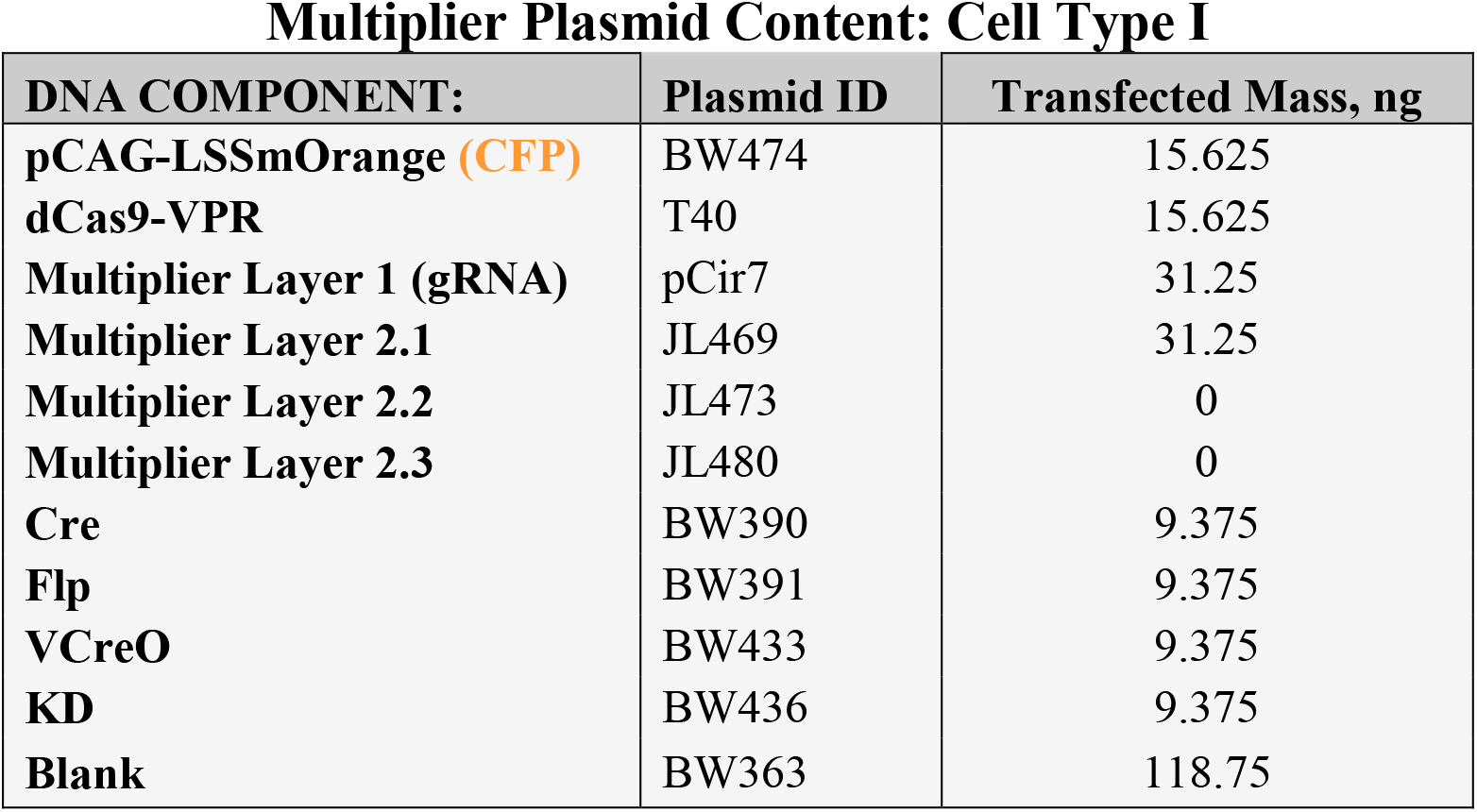

**Table S13:**
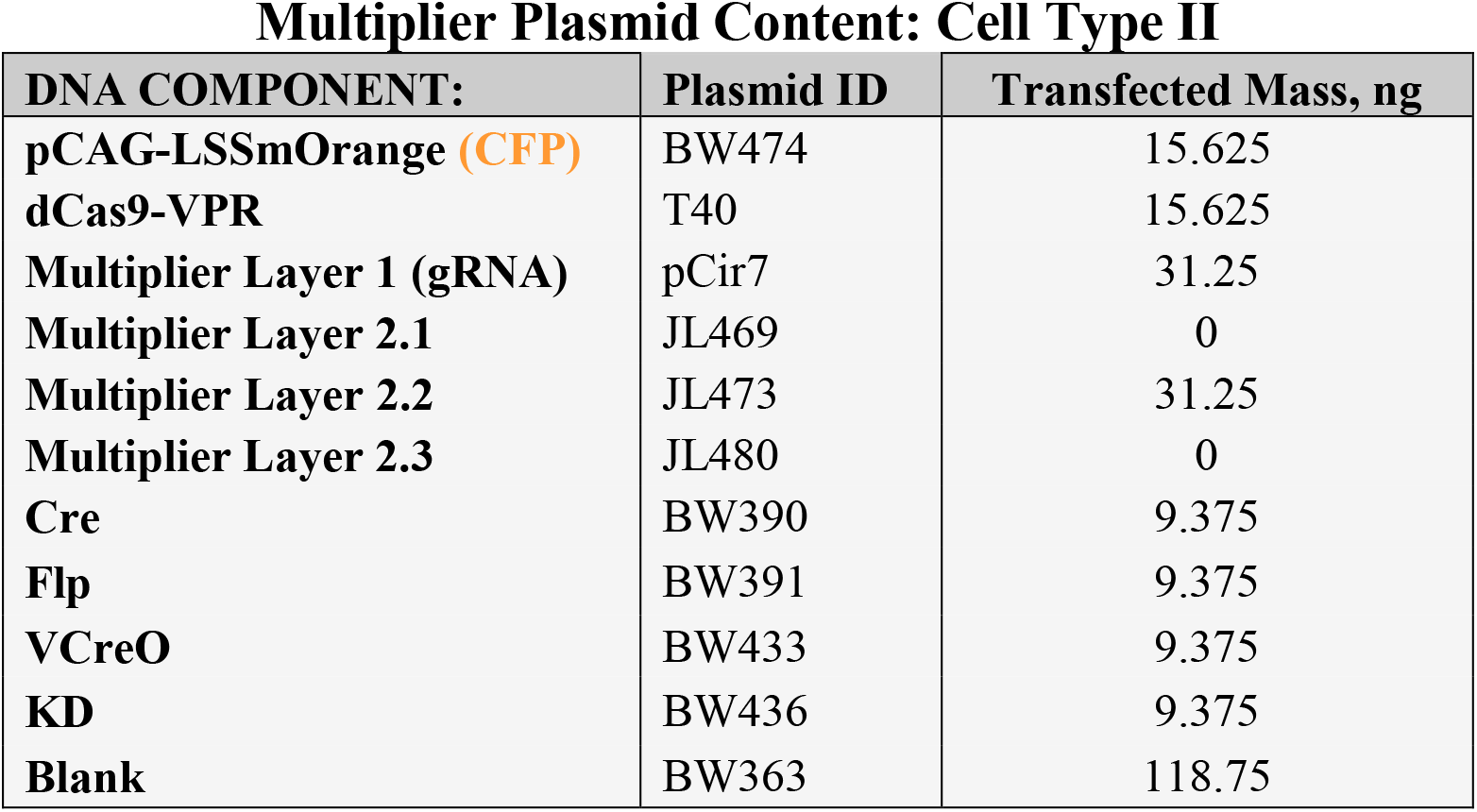
Multiplier Plasmid Content: Cell Type II

| **DNA COMPONENT:** | **Plasmid ID** | **Transfected Mass, ng** |
| --- | --- | --- |
| **pCAG-LSSmOrange (CFP)** | BW474 | 15.625 |
| **dCas9-VPR** | T40 | 15.625 |
| **Multiplier Layer 1 (gRNA)** | pCir7 | 31.25 |
| **Multiplier Layer 2.1** | JL469 | 0 |
| **Multiplier Layer 2.2** | JL473 | 31.25 |
| **Multiplier Layer 2.3** | JL480 | 0 |
| **Cre** | BW390 | 9.375 |
| **Flp** | BW391 | 9.375 |
| **VCreO** | BW433 | 9.375 |
| **KD** | BW436 | 9.375 |
| **Blank** | BW363 | 118.75 |

**Table S14:** Multiplier Plasmid Content: Cell Type III

| **DNA COMPONENT:** | **Plasmid ID** | **Transfected Mass, ng** |
| --- | --- | --- |
| **pCAG-LSSmOrange (CFP)** | BW474 | 15.625 |
| **dCas9-VPR** | T40 | 15.625 |
| **Multiplier Layer 1 (gRNA)** | pCir7 | 31.25 |
| **Multiplier Layer 2.1** | JL469 | 0 |
| **Multiplier Layer 2.2** | JL473 | 0 |
| **Multiplier Layer 2.3** | JL480 | 87.5 |
| **Cre** | BW390 | 9.375 |
| **Flp** | BW391 | 9.375 |
| **VCreO** | BW433 | 9.375 |
| **KD** | BW436 | 9.375 |
| **Blank** | BW363 | 62.5 |

**Table S15:**
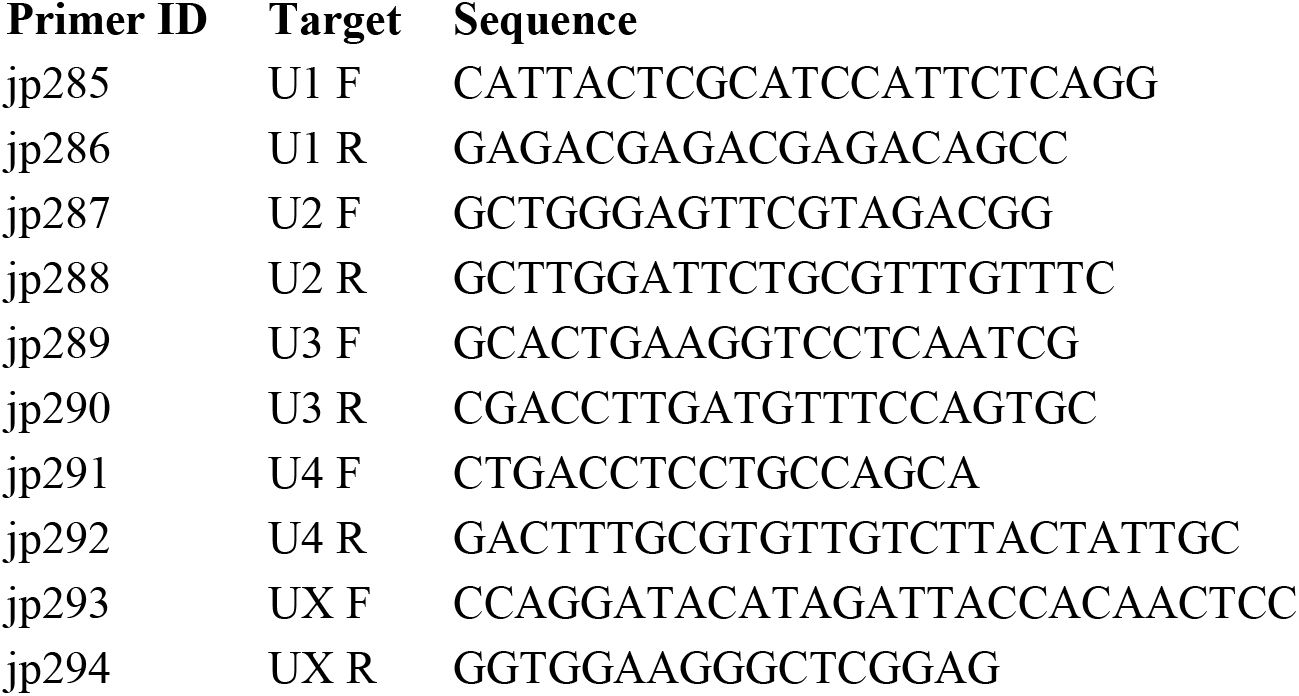

**FIG S1:**
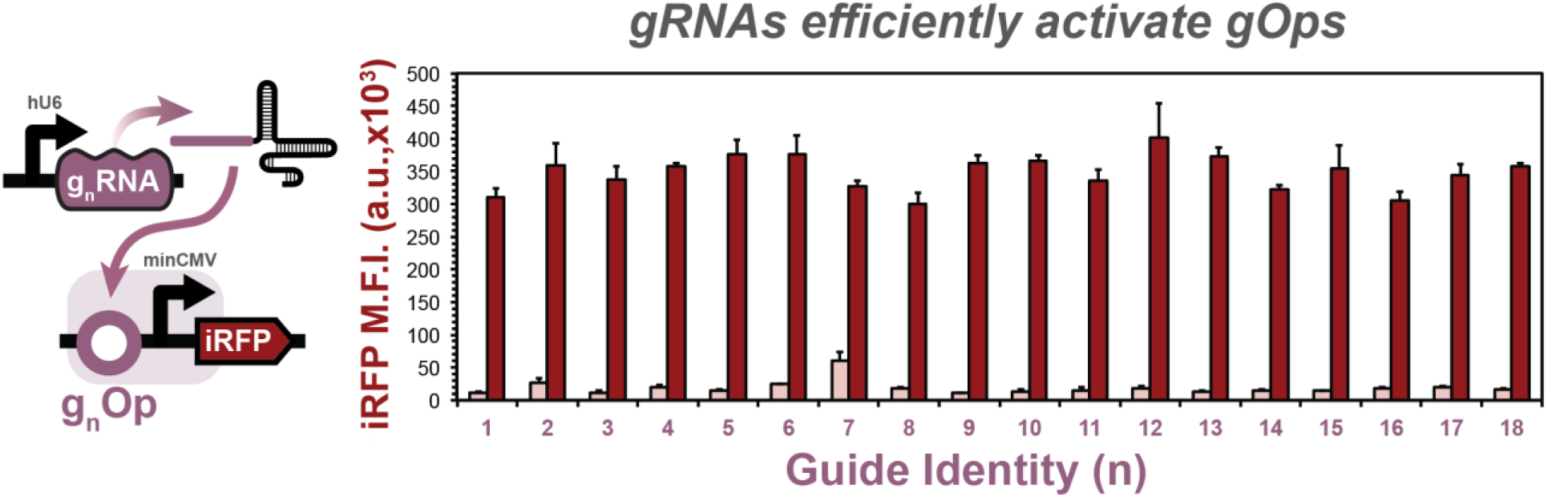
gRNA candidates display high dynamic range, low basal activity. gRNAs targeting consensus operator sites upstream of a minimal CMV (minCMV) promoter were screened for activity in mammalian cells. Each gRNA was transfected with its cognate reporter in the presence (dark red bars) or absence (white bars) of a dCas9-VPR TF. All 18 candidates screened display high dynamic range and low basal promoter activity. Data represent the geometric mean and standard deviation of flow cytometry data collected 48 post transient transfection from three technical replicates of HEK293FT cells (*n = 3*).

**FIG S2:**
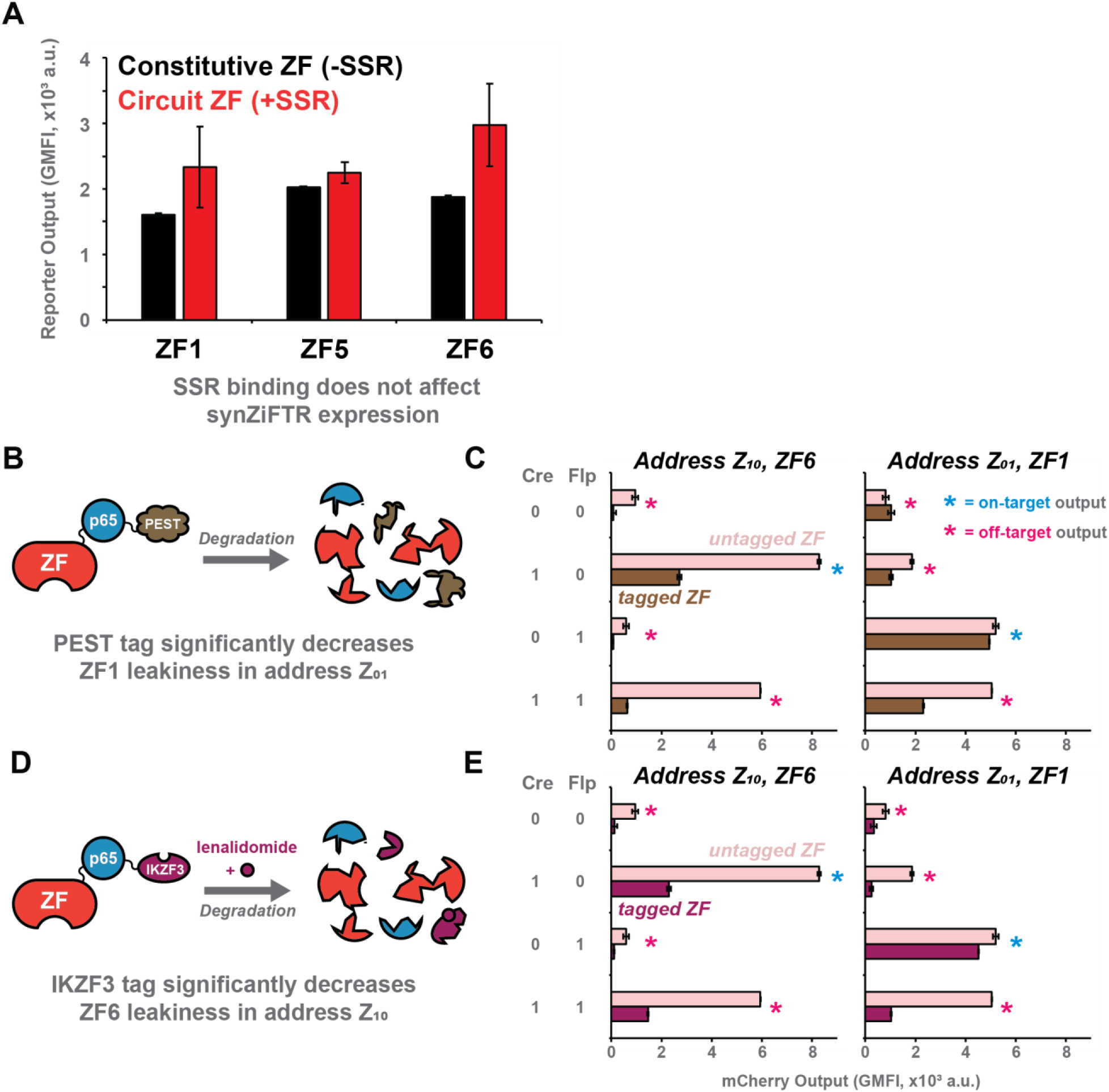
synZiFTR circuits unaffected by SSR target sites, leaky TF expression controlled by degron domain tags. **A**) synZiFTRs expressed from a cleaved Layer 1 circuit that contain 5’ SSR target sites (red bars) display little difference in their ability to drive reporter output signal compared with constitutively expressed synZiFTRs with no 5’ SSR target sites (black bars). Leaky synZiFTR expression from addresses Z10 and Z01 can be mitigated by inducing more rapid turnover of the proteins through PEST (**B-C**) and IKZF3 (**D-E**) degron tags. Data represent the geometric mean and standard deviation of flow cytometry data collected 48 post transient transfection from three technical replicates of HEK293FT cells (n = 3).

**FIG S3:**
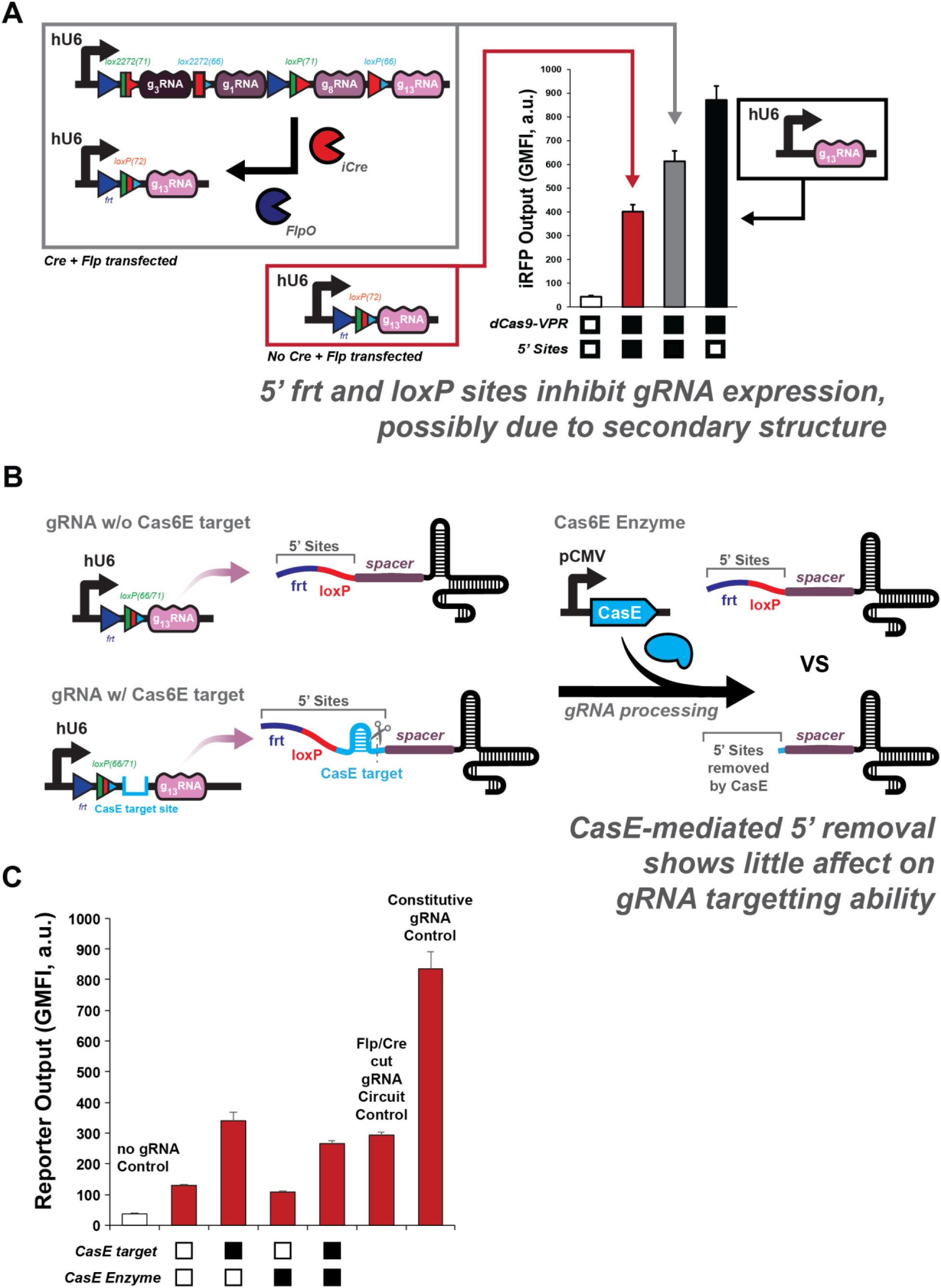
gRNA targeting likely unaffected by additional 5’ nucleotides. **A**) constitutive gRNA expression with no 5’ SSR target sites drives higher expression of reporter output signal than gRNAs containing 5’ SSR target sites, regardless of SSR presence. Attempts to resolve this by tagging gRNA transcripts with CasE target sites (**B**) show little effect (**C**). Data represent the geometric mean and standard deviation of flow cytometry data collected 48 post transient transfection from three technical replicates of HEK293FT cells (n = 3).

**FIG S4:**
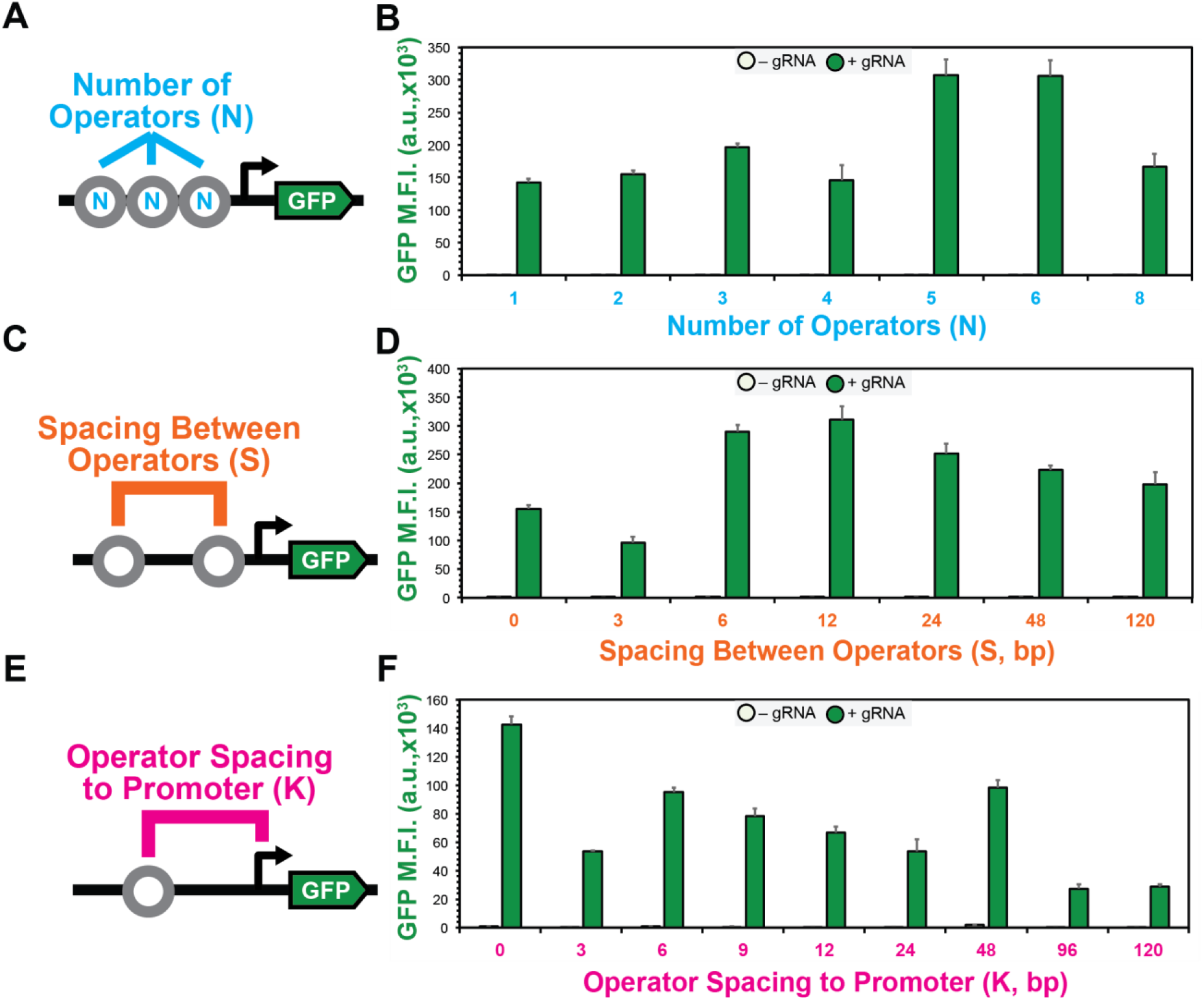
gRNA operator placement affects target promoter transcriptional output strength. gRNA promoters were designed by characterizing the effects of the number of gRNA operator sites (**A-B**), spacing between operator sites (**C-D**) and distance between the operator site and core minimal promoter (minCMV) (**E-F**) on transcriptional output strength. Data represent the geometric mean and standard deviation of flow cytometry data collected 48 post transient transfection from three technical replicates of HEK293FT cells (n = 3).

**FIG S5:**
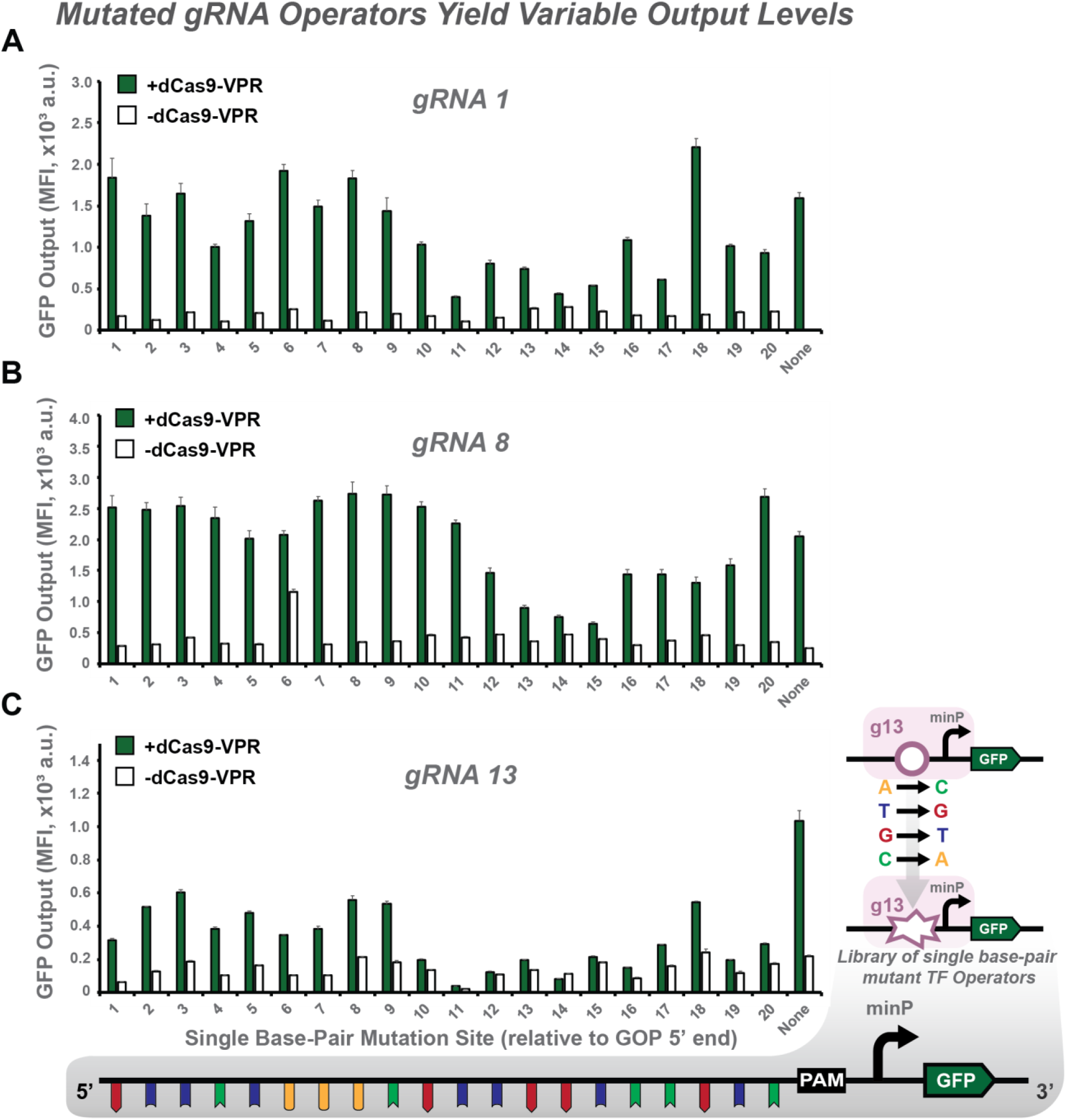
Single base mutations to gRNA operator sites modulates transcription strength of target promoters. To design libraries of gRNA-responsive promoters exhibiting a wide array of transcriptional output strengths, we created single-base mutations in the gRNA operator sites for each gRNA in the Layer 1 circuit. General trends seem to indicate that bases 11-17 from the 5’ end of the operator site are most important for gRNA affinity for its target, observed by the largest fold changes in activity compared with the consensus operator sequence. Data represent the geometric mean and standard deviation of flow cytometry data collected 48 post transient transfection from three technical replicates of HEK293FT cells (n = 3).

**FIG 6:**
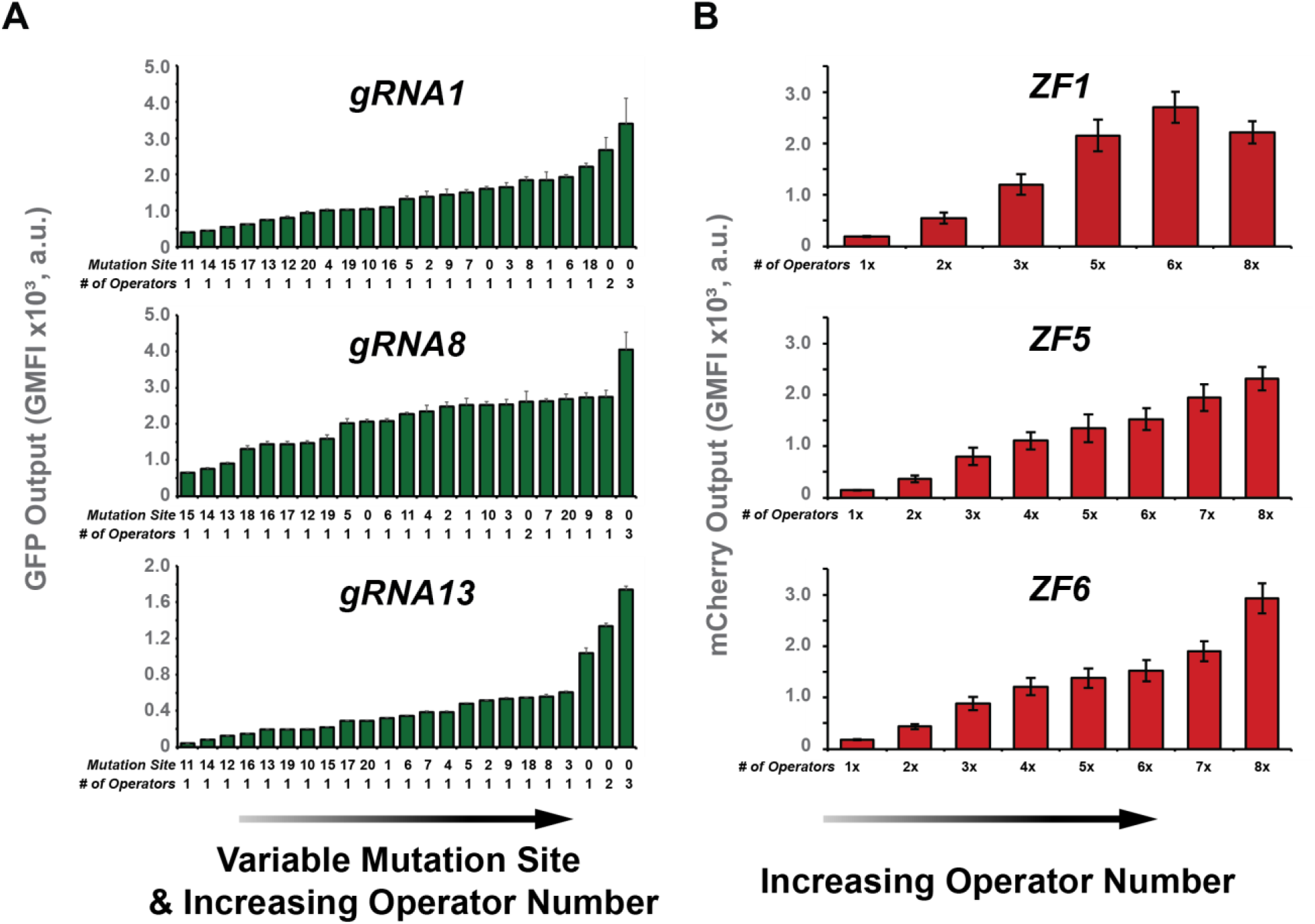
Wide range of transcriptional output available for each programmable TF used in this work. Libraries of gRNA (**A**) and synZiFTR (**B**) responsive promoters demonstrate a wide dynamic range achievable for each system using the CREATE platform. gRNA promoters libraries were constructed by varying the number of TF operator sites and sequence upstream of a minCMV promoter. synZiFTR promoters were designed to include a variable number of TF operator sites upstream of a ybTATA promoter [112]. Data represent the geometric mean and standard deviation of flow cytometry data collected 48 post transient transfection from three technical replicates of HEK293FT cells (n = 3).

**FIG S7:**
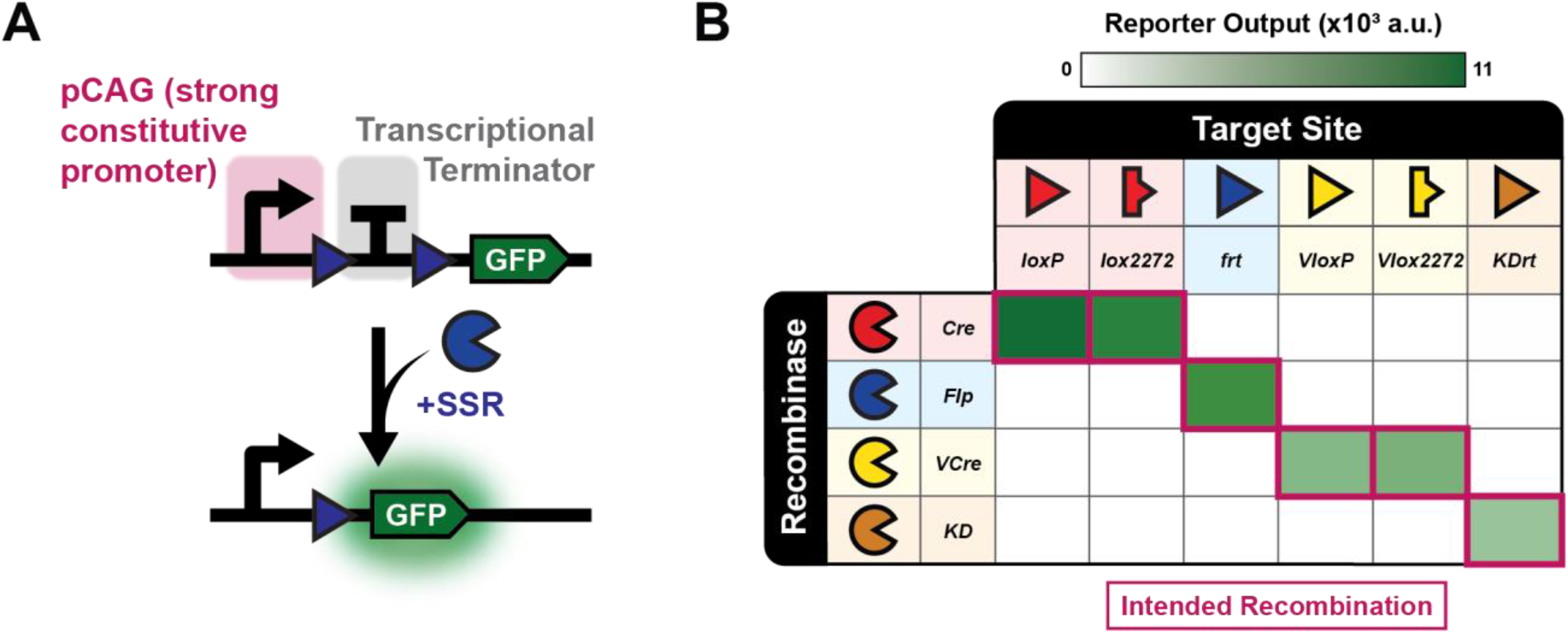
Recombinases and target sites display complete orthogonality, high dynamic range. Cre, Flp, VCre, and KD SSRs were screened against all target sites used in this work by transfecting constitutively expressing SSRs with reporter circuits depicted in (**A**). Each SSR shows complete orthogonality (**B**) against non-cognate target sites while promoting high levels of reporter output from circuits containing the intended target sites. Data represent the geometric mean and standard deviation of flow cytometry data collected 48 post transient transfection from three technical replicates of HEK293FT cells (n = 3).

**FIG S8:**
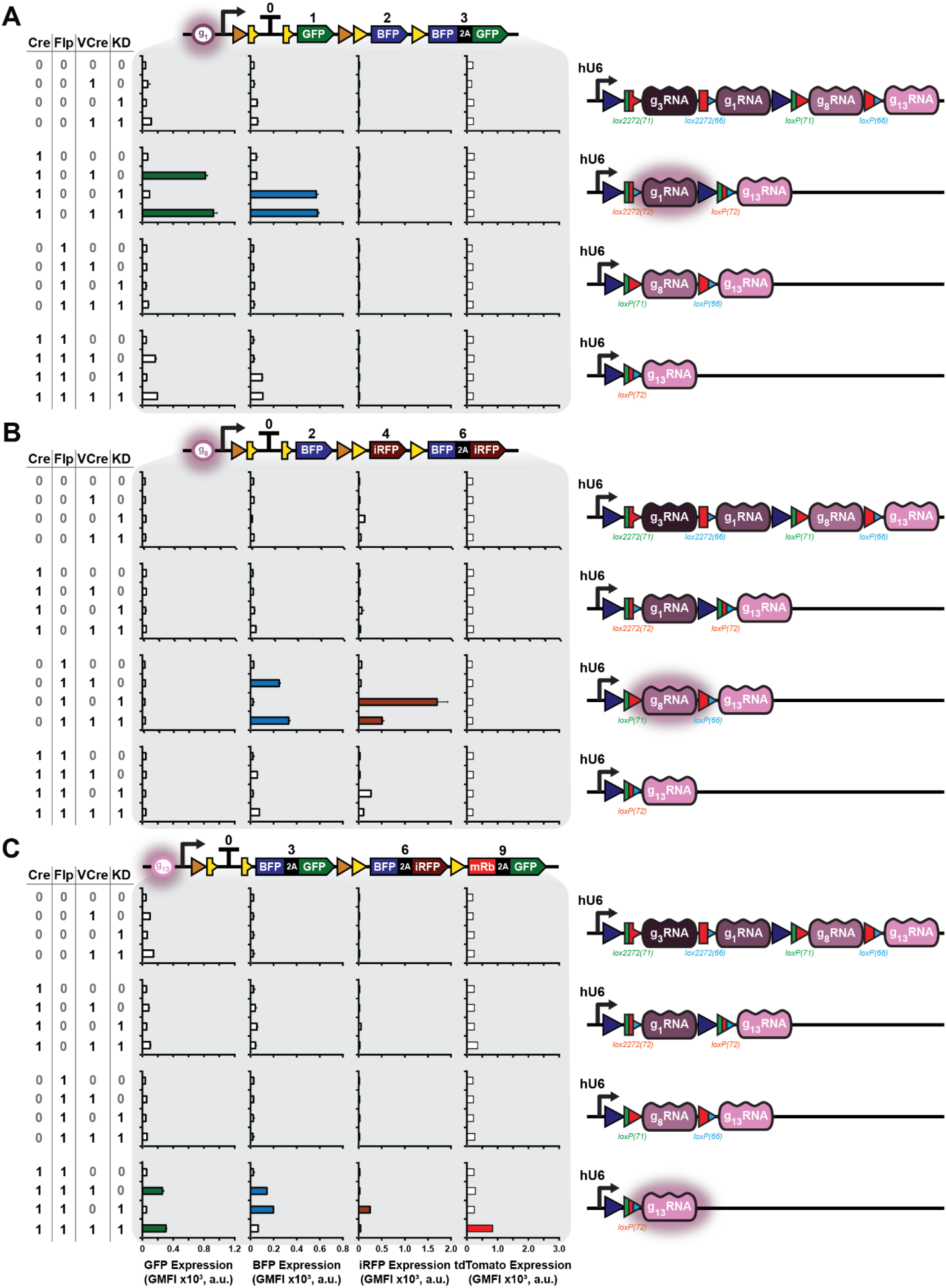
Layer 1 and 2 circuits display orthogonality, some basal activity. Layer 1 circuits were transfected with all possible combinations of SSR inputs and screened for their ability to activate Layer 2 circuits. Some leaky activity from addresses Z10 and Z01 is present, but all gRNAs appear largely orthogonal. Data represent the geometric mean and standard deviation of flow cytometry data collected 48 post transient transfection from three technical replicates of HEK293FT cells (n = 3).

**FIG S9:**
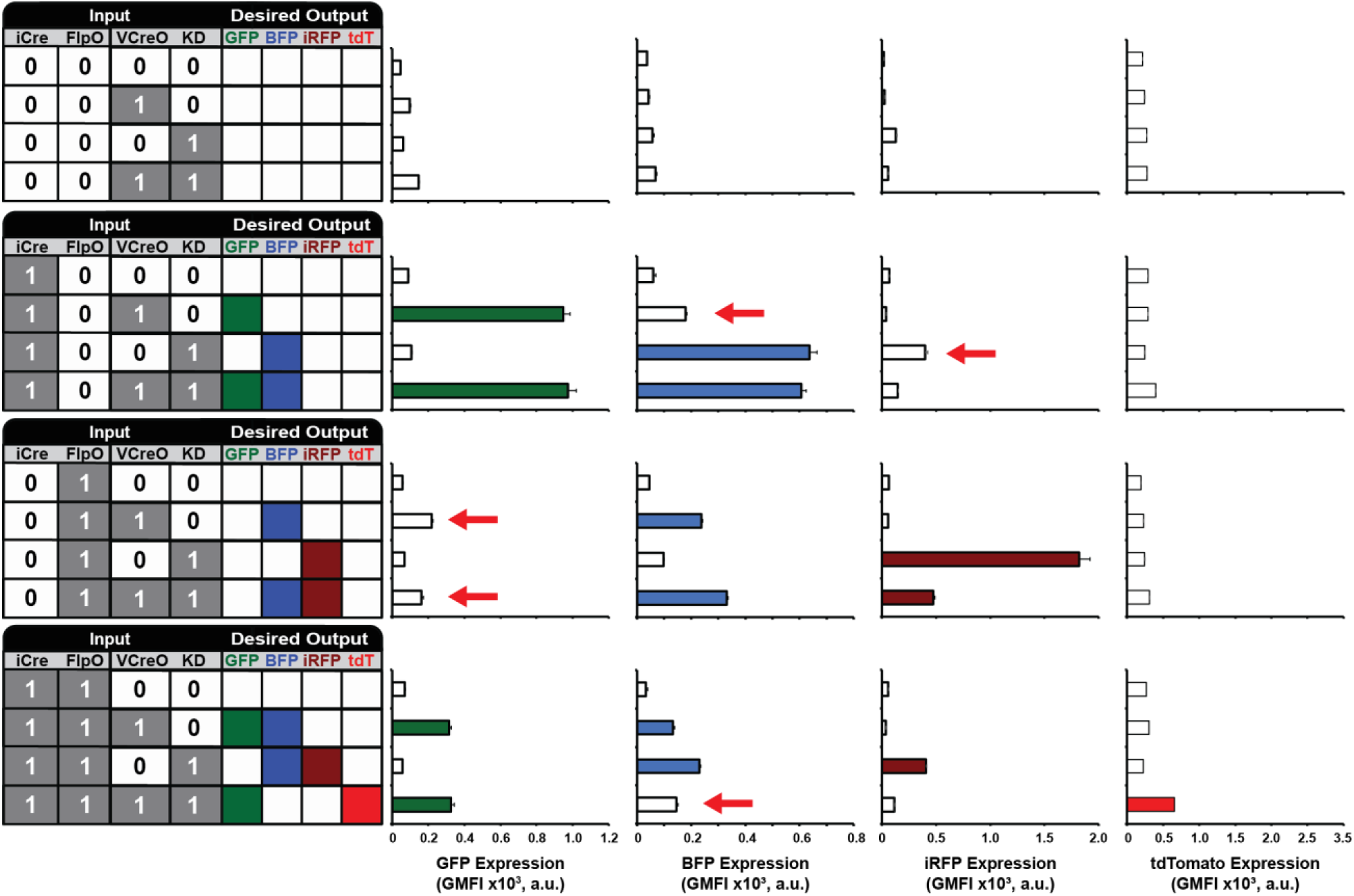
Layer 1 and 2 multiplier circuit plasmids transfected together yield high basal activity in several states. When all Layer 1 and Layer 2 circuits are transfected together in the same cell, basal leaky activity from off-target address activation leads to a breakdown in circuit function and loss of logic-processing capabilities. Red arrows highlight addresses expressing off-target fluorescent protein expression that is indistinguishable from other on-target states. Data represent the geometric mean and standard deviation of flow cytometry data collected 48 post transient transfection from three technical replicates of HEK293FT cells (n = 3).

**FIG S10:**
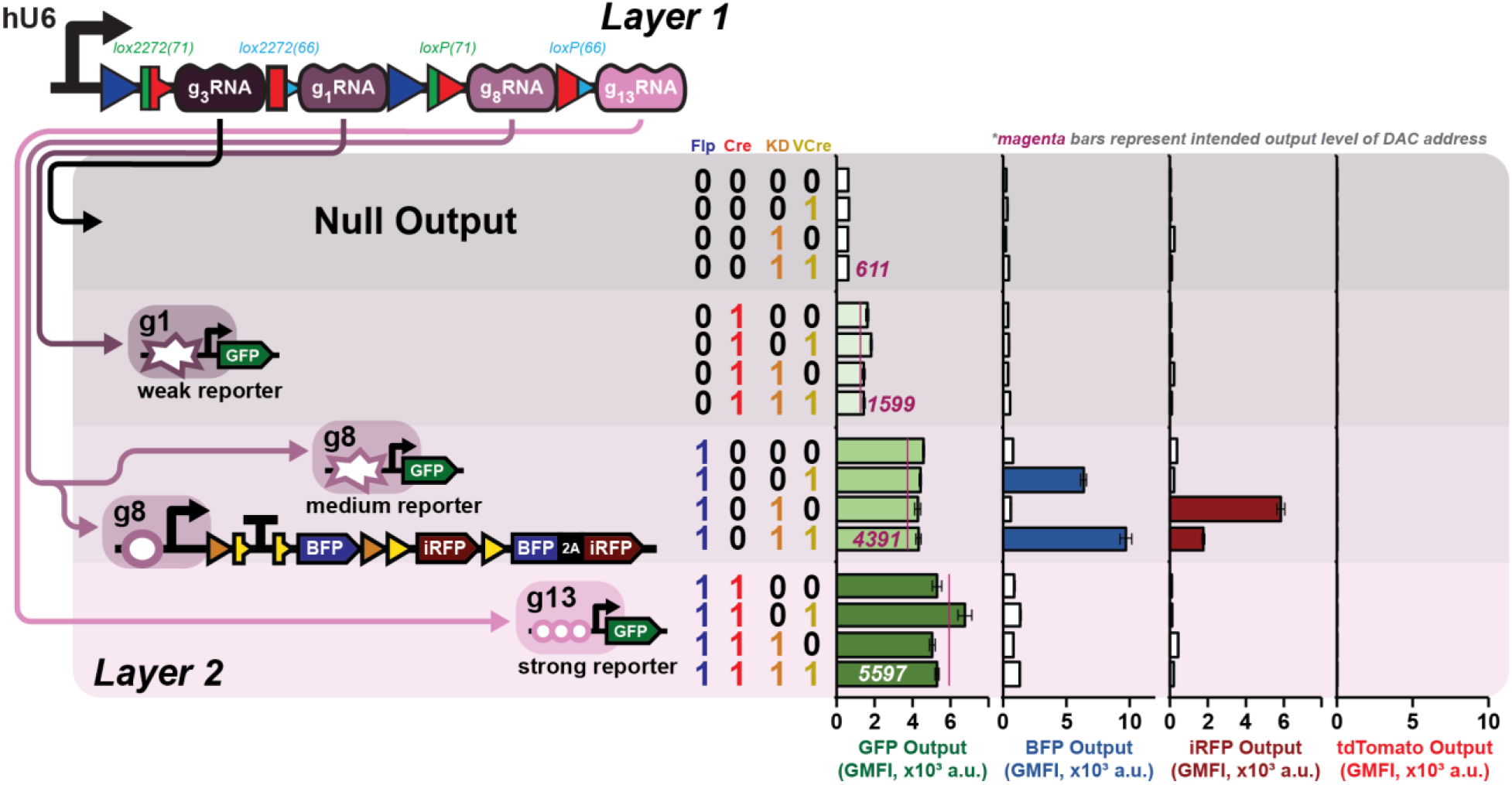
Mixed-signal generator can be composed by simply combining DAC with 2-input-4-output Layer 2 circuit. Transfecting both a ramp up GFP DAC and 2-input-4-output Layer 2 circuit in the same cell with a Layer 1 TF Decoder circuit yields the intended expression trends from both digital and analog signaling components. GFP expression steadily increases as Cre and Flp process the Layer 1 circuit to reveal different TFs, while VCre and KD control the expression of BFP and iRFP reporters from Layer 2 while g8 is expressed from Layer 1. Data represent the geometric mean and standard deviation of flow cytometry data collected 48 post transient transfection from three technical replicates of HEK293FT cells (n = 3).

